# Multivariate autoregressive model estimation for high dimensional intracranial electrophysiological data

**DOI:** 10.1101/2021.12.01.470804

**Authors:** Christopher Endemann, Bryan M. Krause, Kirill V. Nourski, Matthew I. Banks, Barry Van Veen

## Abstract

Fundamental to elucidating the functional organization of the brain is the assessment of causal interactions between different brain regions. Multivariate autoregressive (MVAR) modeling techniques applied to multisite electrophysiological recordings are a promising avenue for identifying such causal links. They estimate the degree to which past activity in one or more brain regions is predictive of another region’s present activity, while simultaneously accounting for the mediating effects of other regions. Including in the model as many mediating variables as possible has the benefit of drastically reducing the odds of detecting spurious causal connectivity. However, effective bounds on the number of MVAR model coefficients that can be estimated reliably from limited data make exploiting the potential of MVAR models challenging. Here, we utilize well-established dimensionality-reduction techniques to fit MVAR models to human intracranial data from ∽100 – 200 recording sites spanning dozens of anatomically and functionally distinct cortical regions. First, we show that high dimensional MVAR models can be successfully estimated from long segments of data and yield plausible connectivity profiles. Next, we use these models to generate synthetic data with known ground-truth connectivity to explore the utility of applying principal component analysis and group least absolute shrinkage and selection operator (LASSO) to reduce the number of parameters (connections) during model fitting to shorter data segments. We show that group LASSO is highly effective for recovering ground truth connectivity in the limited data regime, capturing important features of connectivity for high-dimensional models with as little as 10 s of data. The methods presented here have broad applicability to the analysis of high-dimensional time series data in neuroscience, facilitating the elucidation of the neural basis of sensation, cognition, and arousal.

## Introduction

Measuring causal relationships between activity in different regions of the brain is fundamental to understanding its functional organization. Standard measures of these causal interactions (i.e., effective connectivity) such as Granger Causality (GC) (Granger, 1969; Seth et al., 2015 and generalized partial directed coherence (gPDC) {Baccala, 2007 #9350) can be obtained by fitting linear multivariate autoregressive (MVAR) models to data recorded simultaneously from different locations in the brain. MVAR models estimate the present value measured at a target location using a weighted sum of past values from all other measurement locations. Importantly, because of their multivariate nature, MVAR models represent the interactions between any two locations while simultaneously accounting for the mediating effects of other sites (Blinowska et al., 2004). However, these models are difficult to implement in practice for even modest numbers of recording sites because of the number of parameters that need to be estimated. To fit an MVAR model to a dataset with *M* recording sites and model order (i.e., the number of past values used to estimate present values) of *p*, the number of parameters that must be estimated is equal to *M*^*2*^*p*. The length of stationary data required to reliably estimate MVAR models using ordinary least-squares (OLS) has been estimated previously to be 10*Mp* (Schlogl and Supp, 2006); we show here that this data length is closer to 100*Mp* when *M* is large. The difficulty of reliably estimating MVAR model coefficients from available data appears to have limited previously reported applications in human electrophysiology to <∼30 electrodes (e.g. see (Korzeniewska et al., 2011) for a relatively high-dimensional example; (Brovelli et al., 2004) are more representative and use four and six channel models). The primary contribution of this paper is to explore MVAR model estimation methods that enable modeling of much larger networks from practical data lengths.

Intracranial encephalographic (iEEG) recordings from neurosurgical patients can simultaneously sample neural activity from hundreds of locations in the brain with high spatial and temporal resolution, and thus are a rich source of multivariate data for exploring causal interactions in brain networks. However, exploiting the opportunity afforded by MVAR modeling with hundreds of electrodes is challenging due to limited data availability. For example, with typical iEEG recording parameters of *Mp* = 200 and sampling frequency of 250 Hz, and a model order of *p* = 8, 100*Mp* translates into a required data length of >10 minutes for a single static estimate of connectivity; dynamic connectivity analysis (Preti et al., 2017) in this scenario would be impractical. Furthermore, even when working with longer duration recordings, additional limitations such as the presence of artifacts and nonstationarity can limit the total duration of data available for estimating an accurate MVAR model. These considerations motivate the development of methods to fit MVAR models in the limited data regime. In the past, the standard workaround for this problem has been to estimate MVAR models using smaller subsets of selected recording sites. Unfortunately, this removes potential mediating variables and connections and distorts the true causal network underlying brain activity, drastically increasing the likelihood of detecting spurious effective connections (Granger, 1980; Kus et al., 2004; Olejarczyk et al., 2017).

There have been several attempts to implement dimensionality-reduction techniques to more accurately fit MVAR models to large multivariate datasets. Principal component analysis (PCA) has been used previously to fit MVAR models to scalp EEG data (Joliffe and Morgan, 1992). In this approach, PCA is applied to the electrode-by-electrode covariance matrix to yield “virtual scalp electrodes”, i.e., electrodes projected onto an orthogonalized basis set that more efficiently captures the spatial variability across electrodes. To retain greater spatial information, PCA has also been applied separately to regions of interest (ROIs) on the scalp determined *a priori*, prior to concatenating the full set of principal components across ROIs as input to the MVAR model (Wang et al., 2016). We will refer to this approach as rPCA. Connections between ROIs and their virtual electrodes can then be aggregated and summarized using “block” measures of connectivity (Faes et al., 2012; Faes and Nollo, 2013).

Sparse regression is another common method to reduce the effective number of estimated parameters in MVAR models (Antonacci et al., 2019; Valdes-Sosa et al., 2005). Unlike PCA, which is a processing step implemented prior to model-fitting, sparse regression approaches use a regularizer during the model-fitting step. The Least Absolute Shrinkage and Selection Operator (LASSO) method (Tibshirani, 1996) regularizes the OLS problem with a penalty based on the L1 norm of the MVAR coefficients. This retains essential coefficients, i.e., those explaining the most variance in the training data, while setting smaller coefficients or those with less explanatory power to zero. The group LASSO (gLASSO) approach encourages sparse connections between nodes by shrinking all the coefficients associated with smaller node-node interactions in the model to zero simultaneously (Bolstad et al., 2011). The gLASSO enhances the interpretability of the resulting sparse model since it focuses on sparsity of connections rather than isolated coefficients.

While both rPCA and gLASSO methods have been used separately in previous studies, they have yet to be combined or systematically compared. In this work, we first show that high dimensional iEEG data (117 – 216 electrodes) can be fit successfully using MVAR models provided sufficient data is available. We then explore the performance of different approaches for recovering connectivity profiles in the limited data regime. Physiologically realistic simulated data is created using network models estimated from high dimensional human intracranial EEG (iEEG) data. The simulated network provides ground truth for evaluation of various estimation methods. We use these data to assess how the combination of gLASSO and rPCA methods compares with rPCA-only, gLASSO-only, and standard OLS methods, showcasing the relative performance of each modeling strategy. Specifically, we measure each modeling strategy’s ability to recover known ground-truth regional connectivity, based on a new block gPDC measure that has superior properties to previously used block gPDC measures. We show that gLASSO can capture essential features of connectivity even in very limited data regimes, motivating an illustration of using gLASSO to assess dynamic connectivity. In addition to these simulation experiments, we apply each method to resting-state data derived from high-dimensional iEEG recordings to demonstrate their relative abilities to recover plausible connectivity profiles.

## Methods

### Overview

Our goal was to develop methods for reliably fitting MVAR models to high dimensional neural data. The performance of rPCA and gLASSO estimation methods were objectively evaluated on simulated data generated by physiologically realistic “ground truth” networks. Data-predictive and network connectivity metrics were employed to assess estimation fidelity as a function of data length. The physiologically realistic networks used as ground truth benchmarks in the simulation study were estimated using long segments of resting state iEEG data recorded from neurosurgical patients. The Methods section is divided into five sub-sections: (1) human subjects and iEEG recordings; (2) overview of MVAR models; (3) ground truth network estimation; (4) methods for estimating MVAR models from simulated data, including rPCA, gLASSO, and rPCA-gLASSO; (5) details of the metrics used for assessing estimator performance, including the block gPDC measure we employed to evaluate regional connectivity.

### Human subjects and iEEG recordings

#### Subjects

iEEG data were used to derive ground-truth networks for simulation and to demonstrate that plausible connectivity profiles could be recovered using the approach described here. These data were obtained from ten neurosurgical patients (6 female, ages 21 - 48 years old, median age 34.5 years old; Table 1). The patients had been diagnosed with medically refractory epilepsy and were undergoing chronic invasive iEEG monitoring to identify potentially resectable seizure foci. All human subjects experiments were carried out in accordance with The Code of Ethics of the World Medical Association (Declaration of Helsinki) for experiments involving humans. The research protocols were approved by the University of Iowa Institutional Review Board and the National Institutes of Health. Written informed consent was obtained from all subjects. Research participation did not interfere with acquisition of clinically required data. Subjects could rescind consent at any time without interrupting their clinical evaluation. All subjects were native English speakers, right-handed, and had left hemisphere language dominance, as determined by Wada test. F, female; L, left; M; male; R, right. The hemisphere with predominant electrode coverage is indicated by the prefix of the subject code.

**Table 1.** Subject demographics.

| Subject | Age | Sex | Seizure focus |
| --- | --- | --- | --- |
| L307 | 32 | M | L insula |
| L357 | 36 | M | L medial temporal |
| R369 | 30 | M | R medial temporal |
| R376 | 48 | F | R medial temporal |
| R429 | 32 | F | R anterior and medial temporal |
| R434 | 39 | F | R medial temporal |
| L442 | 33 | F | R temporal (multiple medial and neocortical) |
| L514 | 46 | M | L insula (anterior) |
| R515 | 21 | F | R medial temporal |
| R532 | 42 | F | R ventral frontal (posterior) |

#### iEEG recordings

Recordings were made using subdural and depth electrodes (Ad-Tech Medical, Oak Creek, WI) (Figure 1A; Supplementary Figure 1). After rejecting electrodes that were located in seizure foci, white matter, or outside the brain, or for noise reasons (see below), the median number of recording sites across the 10 subjects was 174 (range 117 – 216; Supplementary Figure 2; Table 2). Subdural arrays consisted of platinum-iridium discs (2.3 mm diameter, 5-10 mm inter-electrode distance), embedded in a silicon membrane. These arrays provided extensive coverage of temporal, frontal, and parietal cortex (Supplementary Figure 1). Depth arrays (8-12 electrodes, 5 mm inter-electrode distance) targeted insular cortex, hippocampus, and amygdala, and additionally provided coverage of the superior temporal plane and superior temporal sulcus. (Note that because of the extensive coverage of auditory cortical structures in the temporal lobe, and adjacent auditory-related cortical structures in parietal and frontal lobes, our scheme for organizing ROIs is auditory-centric.) A subgaleal electrode, placed over the cranial vertex near midline, was used as a reference in all subjects. All electrodes were placed solely on the basis of clinical requirements, as determined by the team of epileptologists and neurosurgeons (Nourski and Howard, 2015).

**Table 2.** Electrode coverage.

| ROI group | ROI | ROI abbreviation | $N_{\text{subjects}}$ | $n_{\text{sites}}$ |
| --- | --- | --- | --- | --- |
| **Auditory core** | Heschl's gyrus, posteromedial | HGPM | 10 | 50 |
| **Auditory non-core** | Heschl's gyrus, anterolateral | HGAL | 9 | 34 |
|  | Planum temporale | PT | 5 | 18 |
|  | Planum polare | PP | 8 | 18 |
|  | Superior temporal gyrus, posterior | STGP | 10 | 109 |
|  | Superior temporal gyrus, middle | STGM | 10 | 55 |
| **Auditory-related** | Superior temporal gyrus, anterior | STGA | 7 | 16 |
|  | Insula, posterior | InsP | 10 | 37 |
|  | Superior temporal sulcus, upper bank | STSU | 6 | 34 |
|  | Superior temporal sulcus, lower bank | STSL | 6 | 24 |
|  | Middle temporal gyrus, posterior | MTGP | 10 | 137 |
|  | Middle temporal gyrus, middle | MTGM | 9 | 89 |
|  | Middle temporal gyrus, anterior | MTGA | 7 | 29 |
|  | Supramarginal gyrus | SMG | 9 | 107 |
|  | Angular gyrus, anterior | AGA | 7 | 46 |
|  | Angular gyrus, posterior | AGP | 5 | 22 |
| **Prefrontal** | Inferior frontal gyrus, pars opercularis | IFGop | 9 | 29 |
|  | Inferior frontal gyrus, pars triangularis | IFGtr | 10 | 63 |
|  | Inferior frontal gyrus, pars orbitalis | IFGor | 7 | 17 |
|  | Middle frontal gyrus | MFG | 9 | 102 |
|  | Superior frontal gyrus | SFG | 2 | 2 |
|  | Anterior cingulate cortex | ACC | 5 | 5 |
|  | Orbital gyrus | OG | 9 | 103 |
|  | Transverse frontopolar gyrus | TFG | 3 | 10 |
| **Sensorimotor** | Precentral gyrus | PreCG | 9 | 84 |
|  | Postcentral gyrus | PostCG | 9 | 54 |
|  | Paracentral lobule | ParaCL | 1 | 2 |
| **Other** | Premotor cortex | PMC | 10 | 37 |
|  | Cingulate gyrus | CingG | 4 | 25 |
|  | Parahippocampal gyrus | PHG | 7 | 27 |
|  | Fusiform gyrus | FG | 10 | 39 |
|  | Inferior temporal gyrus, posterior | ITGP | 6 | 17 |
|  | Inferior temporal gyrus, middle | ITGM | 8 | 28 |
|  | Inferior temporal gyrus, anterior | ITGA | 8 | 34 |
|  | Temporal pole | TP | 8 | 70 |
|  | Insula, anterior | InsA | 10 | 28 |
|  | Frontal operculum | fOperc | 3 | 6 |
|  | Parietal operculum | pOperc | 2 | 4 |
|  | Gyrus rectus | GR | 5 | 11 |
|  | Superior parietal lobule | SPL | 2 | 8 |
|  | Precuneus | PreCun | 1 | 1 |
|  | Lingual gyrus | LingG | 2 | 4 |
|  | Middle occipital gyrus | MOG | 4 | 10 |
|  | Inferior occipital gyrus | IOG | 1 | 1 |
|  | Amygdala | Amyg | 7 | 19 |
|  | Hippocampus | Hipp | 5 | 16 |
|  | Putamen | Put | 3 | 6 |
|  | Caudate nucleus | Caud | 1 | 1 |
|  | Substantia innominata | SubInn | 1 | 2 |
|  | Ventral striatum | vStr | 1 | 2 |
| **Total** |  |  | 10 | 1692 |

**Figure 1.**
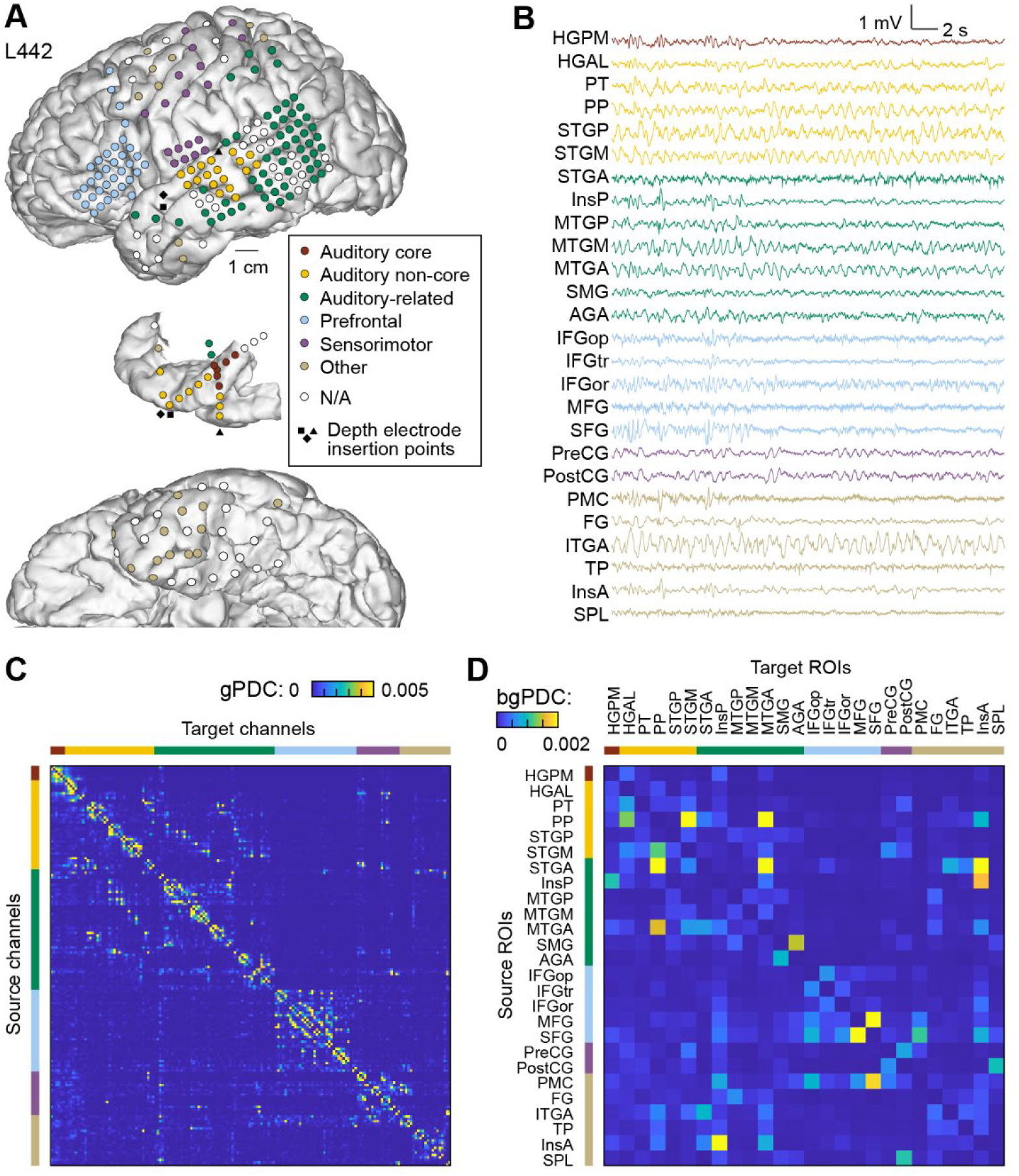
Raw data and ground truth networks. **A**. Example of electrode coverage of the left hemisphere in a representative subject (L442). Top-to-bottom: Lateral view of the left hemisphere, top-down view of the superior temporal plane, ventral view of the left hemisphere. Recording sites are color-coded according to the ROI group. Sites identified as seizure foci or characterized by excessive noise, and depth electrode contacts localized to the white matter are denoted by white symbols. Insertion points of depth electrodes implanted in the superior temporal plane are shown on the lateral view as black symbols. **B**. Example raw data from the subject in A. Data is shown for one recording site in each ROI. **C**. Connectivity for the subject in A estimated from 4 minutes of data fit via a ridge-regression model, =10^−4^. Scalar (electrode level) broad-band gPDC shown. **D**. Block (ROI) connectivity (bgPDC; Eq. 18) for the same subject. Color bars in panels C and D represent ROI groups, color-coded as shown in the legend of panel A. See Table 2 for list of ROI abbreviations.

No-task, resting-state (RS) data were recorded in the dedicated, electrically shielded suite in The University of Iowa Clinical Research Unit while the subjects lay in the hospital bed. Resting state data were collected a median of 5.5 days [range 2 – 11 days] after electrode implantation surgery. In the first two subjects (L307 and L357), data were recorded using a TDT RZ2 real-time processor (Tucker-Davis Technologies, Alachua, FL). In the remaining 8 subjects (R369 through R532), data acquisition was performed using a Neuralynx Atlas System (Neuralynx Inc., Bozeman, MT). Recorded data were amplified, filtered (0.1–500 Hz bandpass, 5 dB/octave rolloff for TDT-recorded data; 0.7–800 Hz bandpass, 12 dB/octave rolloff for Neuralynx-recorded data) and digitized at a sampling rate of 2034.5 Hz (TDT) or 2000 Hz (Neuralynx). For 8/10 subjects, the duration of recordings was 10 mins. For the other two subjects (429R, 532R), the duration was 11 minutes.

### iEEG data analysis

#### Anatomical reconstruction and ROI parcellation

Electrode localization relied on post-implantation T1-weighted structural MR images and post-implantation CT images. All images were initially aligned with pre-operative T1 images using linear coregistration implemented in the FMRIB Software Library (FSL; FLIRT) (Jenkinson et al., 2002). Electrodes were identified in the post-implantation MRI as magnetic susceptibility artifacts and in the CT as metallic hyperdensities. Electrode locations were further refined within the space of the pre-operative MRI using three-dimensional non-linear thin-plate spline warping (Rohr et al., 2001), which corrected for post-operative brain shift and distortion. The warping was constrained with 50-100 control points, manually selected throughout the brain, which aligned to visibly corresponding landmarks in the pre- and post-implantation MRIs.

Regional connectivity was assessed by grouping electrodes based on location. Electrodes were assigned to one of 50 ROIs organized into 6 ROI groups (Figure 1A; Supplementary Figure 1; Supplementary Table 1) based upon anatomical reconstructions of electrode locations in each subject. For subdural arrays, it was informed by automated parcellation of cortical gyri (Destrieux et al., 2010; Destrieux et al., 2017) as implemented in the FreeSurfer software package. For depth arrays, ROI assignment was informed by MRI sections along sagittal, coronal, and axial planes. Heschl’s gyrus (HG) was subdivided into the posteromedial (HGPM) and anterolateral (HGAL) portions (core auditory cortex and adjacent non-core areas, respectively). This division was made using physiological criteria (characteristic short-latency evoked responses to click trains and frequency-following responses in HGPM but not HGAL; see (Brugge et al., 2009) and (Nourski et al., 2016). Superior temporal gyrus (STG) was subdivided into posterior and middle non-core auditory cortex ROIs (STGP and STGM), and auditory-related anterior ROI (STGA) using the transverse temporal sulcus and ascending ramus of the Sylvian fissure as macroanatomical boundaries. Middle and inferior temporal gyrus (MTG and ITG) were each divided into posterior, middle, and anterior ROI by diving the gyrus into three approximately equal thirds along its length. The insula was subdivided into posterior and anterior ROIs, with the former considered within the auditory-related ROI group (Zhang et al., 2019). Within cingulate gyrus, anterior cingulate cortex (as identified by automatic parcellation in FreeSurfer) was considered a prefrontal ROI. Angular gyrus (AG) was divided into posterior and anterior ROIs (AGP and AGA) using the angular sulcus as a macroanatomical boundary. Recording sites identified as seizure foci or characterized by excessive noise, and depth electrode contacts localized to the white matter or outside brain, were excluded from analyses and are not listed in Supplementary Table 1.

#### Preprocessing of iEEG data

For each subject, iEEG data were downsampled to 250 Hz and divided into segments of varying length (10 – 960 s). Artifact rejection involved three steps. First, outlier electrodes were identified based on the average log amplitude in one-minute segments across seven frequency bands computed using the demodulated band transform (DBT; (Kovach and Gander, 2016): delta (1-4 Hz), theta (4-8 Hz), alpha (8-14 Hz), beta (14-30 Hz), gamma (30-50 Hz), high gamma (70-110 Hz), and total (sum of all others). Analytical amplitude measured in each band was *z*-scored across electrodes in each segment and averaged across segments. Electrodes with a mean *z*-score > 3.5 in any band were removed, including from further artifact rejection methods.

Second, intervals containing artifacts in the raw voltage traces were rejected on every electrode. We identified times when any electrode had extreme absolute raw voltage >10 SD for that electrode and marked as artifact the surrounding time until that electrode returned to zero voltage, plus an additional 100 ms before and after. Note that because we were measuring connectivity, any data interval identified on a single electrode was excluded for all electrodes. For each subject, if this procedure identified >1% of recording time as artifact, we optimized the total data kept for that subject (= electrodes × non-artifact time) by further excluding electrodes if retaining those electrodes caused more loss of data on the remaining electrodes via artifact rejection than they themselves contributed.

Third, we applied a specific additional noise criterion to eliminate brief power spikes in the high gamma band, a band that in some subjects was particularly sensitive to noise in our recording environment. We excluded all intervals containing segments in which high gamma power averaged across electrodes was greater than five standard deviations after excluding electrodes and times already removed in steps 1 and 2.

### Multivariate autoregressive models

Multivariate autoregressive (MVAR) models represent the present value at each electrode as a weighted combination of past values at all other electrodes. Let *y*^*m*^*(n)* be the voltage in electrode *m* at time *n* The MVAR model is for *y*^*m*^*(n)* is

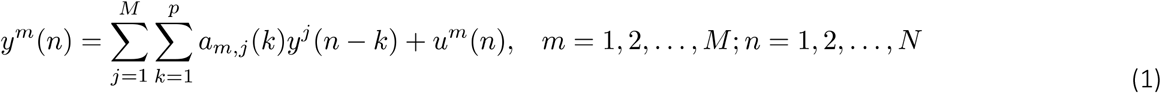

where *a*_*m,j*_*(k)* is the weight applied to electrode *j* at lag *k* to predict electrode *m, u*^*m*^(*n*) is the model error or innovations process in electrode *m, p* is the memory or maximum lag considered by the model, *N* is the number of data samples, and *M* is the number of electrodes. The innovations are typically assumed to be white noise that is uncorrelated for different electrodes. Note that although MVAR models are linear, they can detect both linear and nonlinear causal interactions and may be more robust to noise than methods that explicitly capture nonlinearities (Astolfi et al., 2009; Freiwald et al., 1999; Gourévitch et al., 2006; Korzeniewska et al., 2011; Netoff et al., 2004).

In the large data regime (*N* large) the MVAR model coefficients *a*_*m,j*_*(k)* may be estimated from data using the OLS method (Lutkepohl, 2005). Define

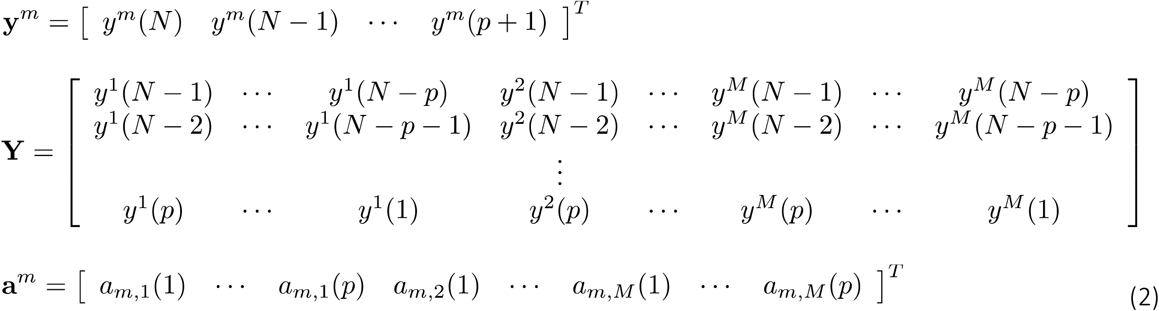

so that the least squares problem for estimating **a**^*m*^ is written in matrix form as

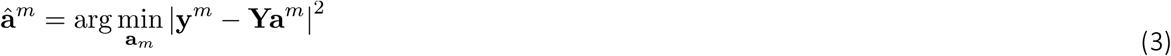

The solution to Eq. (3) is given by

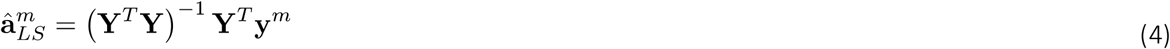

assuming ***Y***^*T*^ ***Y*** is an invertible matrix, which requires *N-p ≥ Mp*.

In practice, the number of stationary data samples must be significantly larger than the minimum to avoid large errors in the MVAR parameter estimates. A rule of thumb of *N > 10Mp* has been suggested (see, e.g., (Schlogl and Supp, 2006)). It is important to note that such guidelines assume the data samples have some degree of statistical independence. Thus, the effective number of samples available is determined by the bandwidth of the signal, not the absolute sampling rate. Oversampling the data does not improve estimation quality. For example, if the data has a bandwidth of 50 Hz, then the number of effective samples available for estimation is approximately 100 samples/s. In our data set we had several subjects with more than 200 electrodes. If we assume a modest memory of *p = 8*,, then the *N > 10Mp* rule of thumb suggests we need at least 16,000 statistically independent samples of data. If, for example, we assume an approximate 50 Hz bandwidth, then 16,000 samples map to 160 s or nearly three minutes of data. The strong 1/f characteristic of EEG data may reduce the effective bandwidth of the data in many situations, especially when strong lower frequency rhythmic activity is present, and thus significantly more data may be required in some situations.

The requirement of long, stationary data segments significantly limits the utility of the OLS approach to MVAR model estimation for large networks and is the primary motivation for the evaluation of the rPCA and gLASSO approaches described below.

### Ground-truth network estimation

The “true” networks associated with our human subject data were unknown, which makes it impossible to evaluate objectively the performance of any MVAR model estimation algorithm from measured data. Hence, we used our human subject data to obtain physiologically realistic ground-truth MVAR models. The ground truth models were then used to generate simulated data for objective evaluation of the performance of MVAR model estimation algorithms as a function of data duration.

Exploration of our data revealed that the OLS solution for the MVAR model coefficients gave spurious results in some electrodes even with data record lengths exceeding four minutes, so we employed ridge regression (Hoerl and Kennard, 2000) to regularize the estimates for the ground-truth models. That is, instead of Eq. (3), we chose the ground truth models according to

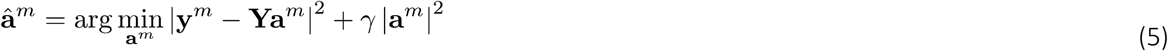

where the regularization parameter γ is chosen as

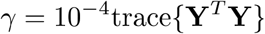

This value for γ was chosen empirically to provide sufficient regularization in electrodes that had spurious connectivity in the OLS approach without significantly altering the connectivity in the remaining electrodes. The solution to Eq. (5) is given by

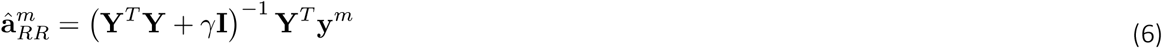

We chose a model order *p =* 8 for all ground-truth models for several reasons. First, initial evaluation suggested that the quality of the model predictive performance did not improve appreciably using larger values of *p*. Second, we wanted the physiologically inspired ground-truth models - derived from different subjects - to use the same model order to facilitate evaluation of algorithm performance across different ground-truth models. Commonly used methods such as Akaike Information Criterion or cross-validation give different optimal values of *p* for different subjects. Third, the data required for model estimation is proportional to *p*, so we chose to work with smallest plausible value. Fourth, and perhaps most importantly, the main goal of this process is to create physiologically inspired models for the simulation study. Achieving this goal does not require precise determination of model order.

One ground-truth network model was created for each subject for a total of ten different models.

Simulated data was created from the network model by applying white noise input as the innovations process *u*^*m*^ (*n*) with variance 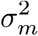 given by the estimated prediction error variance

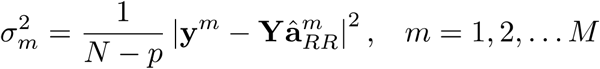

where **y**^*m*^ and **Y** are defined based on thirty s of test data after the four minutes used to estimate 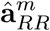 from each subject.

### Model estimation methods

Simulated data from the ten ground-truth models was used to evaluate the performance of four different model MVAR estimation methods as a function of data record length: 1) OLS (Eq. (3)); ROI-based principal component analysis (rPCA) with OLS; 3) Self-connected group LASSO (gLASSO); and 4) rPCA with gLASSO (rPCA-gLASSO). The rPCA, gLASSO, and rPCA-gLASSO methods are described next.

#### ROI-Based Principal Component Analysis (rPCA)

One of the approaches we employed to reduce model dimension – and consequently data requirements – was applying PCA to the collection of electrode signals associated with each ROI. Let **Y**_*j*_*(n)* be an *M*_*j*_*-by-1* vector containing the *M*_*j*_ electrodes of data associated with the *M*_*j*_ electrodes in the *j*^*th*^ ROI. We map **y**_*j*_*(n)* into an *L*_*j*_*-by-1* (*L*_*j*_ *≤ M*_*j*_)vector of PCA signals **x**_j_ *(n)* as

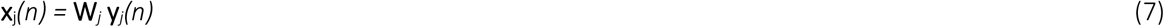

where **W**_*j*_ is an *L*_*j*_*-by-M*_*j*_ matrix whose rows are the eigenvectors corresponding to the largest eigenvalues of the covariance matrix of **y**_*j*_*(n)*. The number of PCA components *L*_*j*_ was chosen so that **x**_*j*_*(n)* represents a specified fraction of the variance associated with **y**_*j*_*(n)*. We chose *L*_*j*_ to represent 95% variance in the results shown later. In our data, these thresholds reduced the total number of electrodes by a factor of approximately two. Note that given PCA signals **x**_*j*_*(n)* and **W***j* one can project back to identify the corresponding approximated electrode data as

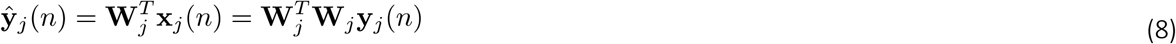

rPCA is an aggregation approach that creates virtual electrodes representing the unique components in the ROI. This approach to reducing the dimensionality of the network is depicted in Figure 2, which also illustrates a segment of the original data signals **y**_*j*_ (*n*), the virtual (PCA) signals **X**_*j*_ (*n*), and the back-projected data signals **ŷ**_*j*_ (*n*) from HGAL in one subject. While the connectivity between the original physical electrodes is modified when operating in the rPCA space, the connectivity between ROIs is preserved to the extent that all significant components of the electrode data **y**_*j*_*(n)* are retained. It is straightforward to show that if 100% of the variance is retained in all regions, then **W**_*j*_ is an invertible matrix and the connectivity between ROIs is identical between electrode and PCA spaces (see Supplementary Methods). As the fractional variance retained decreases, model complexity decreases, resulting in reduced computation and data requirements, but the potential for distortion of connectivity between ROIs increases.

**Figure 2.**
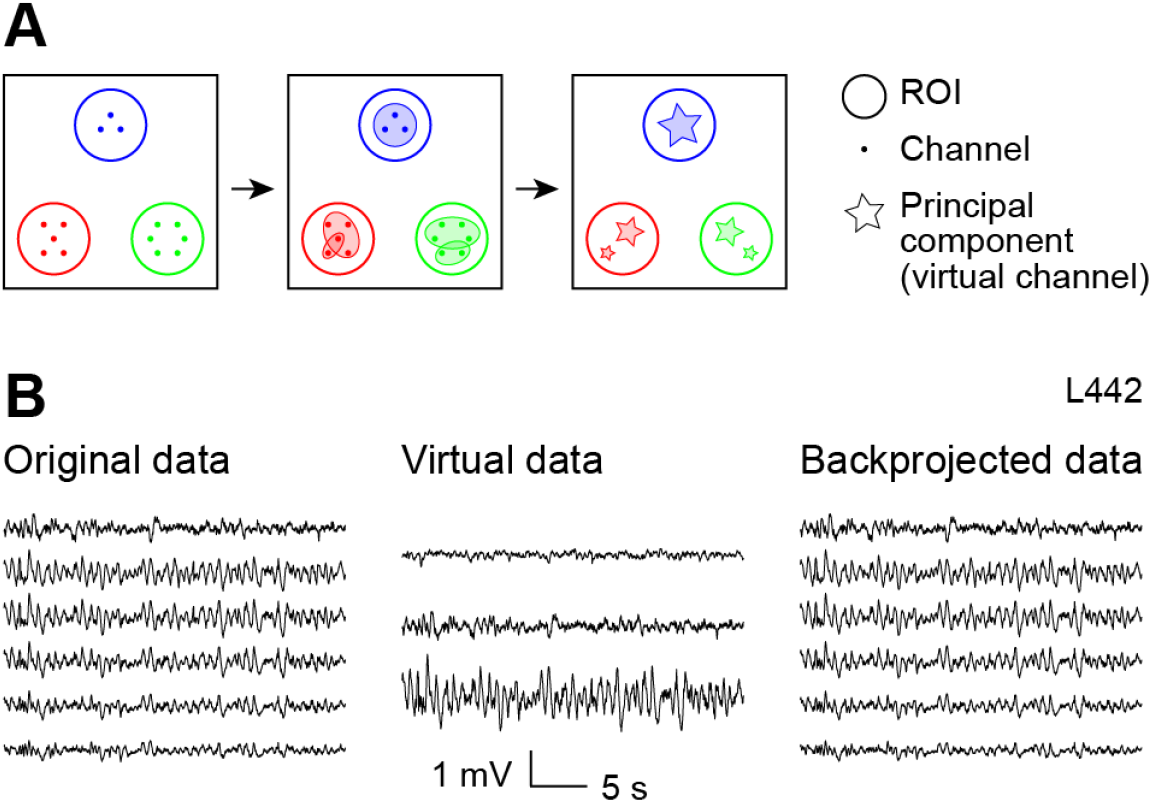
Illustration of rPCA method. A: PCA was applied to electrodes within pre-assigned ROIs (see Figure 1) to create virtual electrodes. B: Illustration of rPCA applied to one ROI (HGAL) in one subject, showing original data (*left*), virtual electrode representation of these data (*middle*), and data backprojected into original data space (*right*).

The MVAR model for the rPCA data is obtained by rewriting Eq. (1) as

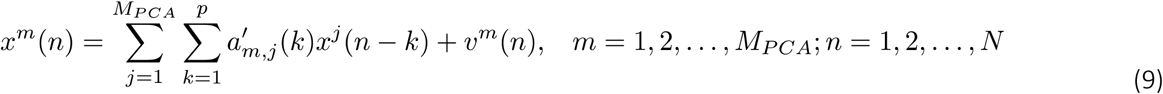

where 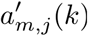 is the rPCA MVAR model coefficient representing the influence of PCA electrode *j* at lag *k* on PCA electrode *m*, υ^*m*^ (*n*) is the innovation or model error in PCA electrode *m*, and *M* _*P C A*_ is the number of virtual electrodes in the model. OLS estimates for the rPCA MVAR model coefficients are obtained analogously to Eqs. (3) and (4).

#### Self-Connected Group LASSO (gLASSO)

The self-connected gLASSO (Bolstad et al., 2011) was evaluated to determine the reduction in data requirements possible using a penalty that encouraged sparse connectivity between physical or virtual electrodes. Define a *p*-by-1 vector of coefficients associated with the connection from electrodes *j* to *m*

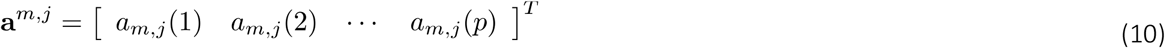

so that **a**^*m*^ in Eq. (2) is expressed as

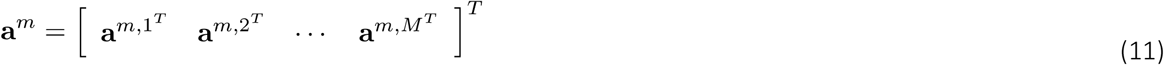

The gLASSO problem is then expressed as

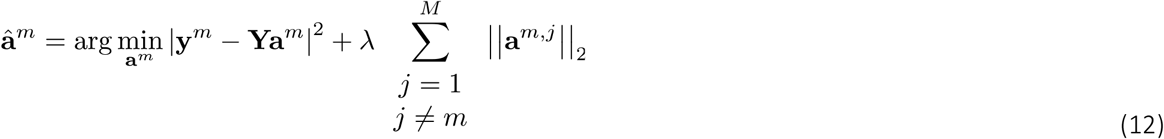

Use of the two-norm of **a**^*m,j*^ in the regularization term ensured that coefficients relating electrode *j* to electrode *m* were penalized as a group, that is, they were set to zero as a group. Note that self-connections were not penalized, which is why *j=m* was excluded from the regularizer. Larger values of the regularization parameter λ resulted in a smaller number of nonzero connections at the expense of larger modeling error.

The solution to Eq. (12) was obtained using standard convex optimization methods based on the code of (Bolstad et al., 2011). Five-fold cross validation was used to select λ:

1. Split available data into five equal portions or folds.
2. Use all folds but the *k*^th^ fold of data to train different models for a range of 11 candidate λ values. The candidate values for λ were chosen by first finding the smallest λ which guarantees all coefficients are zero (Bolstad et al., 2011). The 11 candidate values were chosen evenly spaced between 0% (OLS) and 40% of this value. This maximum was chosen based on the observation that the models have very few nonzero connections at and beyond 40% of λ that guarantees all zero connections.
3. Use the *k*^th^ fold to determine the squared prediction error of each candidate model when applied to data not used to train the model.
4. Repeat steps 2 and 3 five times with different folds held out from training and average the resulting squared errors.
5. Choose the model corresponding to the value λ that results in the minimum averaged error.

This process was repeated for each target electrode *m* in the model. Thus, the coefficient vectors used to predict different electrodes may have had different levels of sparsity. Some areas of the brain are more densely connected than others, so it is plausible that the connectivity to different electrode locations has varying levels of sparsity.

Note that the gLASSO regularizer in Eq. (12) is known to shrink the model coefficients in a manner that depends on λ (Bolstad et al., 2011). Hence, after determining which connections were present using the gLASSO procedure, we estimated the coefficients associated with the active connections by solving a least-squares problem involving only those connections. That is, the gLASSO was used for subset selection — to determine which subset of the connections should be included in the model – and least squares was used to estimate the corresponding coefficients of the reduced complexity model.

The acronym rPCA-gLASSO refers to the process of applying the gLASSO method to the rPCA model described in Eq. (9).

### Performance characterization

The performance of four different modeling approaches was evaluated using simulated data. We simulated the specified length of data for each of ten subjects using the corresponding ground truth model and estimate model coefficients from the simulated data. All estimated MVAR models assumed the correct model order of *p =* 8 so that we could evaluate estimation algorithm performance independent of model order selection. This process was repeated ten times for each ground truth model and data length and the performance averaged over these ten trials. Two different measures were employed to characterize model performance: the model prediction error on data not used to train the model and comparison between the connectivity estimated from the estimated MVAR model coefficients and the ground-truth network connectivity of the MVAR process used to create the simulated data.

It is useful to rewrite Eq. (1) in matrix form as

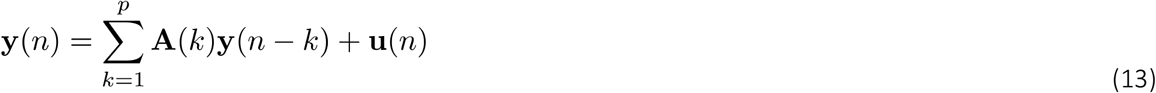

where **y***(n)* is the *M-by-1* vector of data at all *M* electrodes, **A***(k)* is an *M-by-M* matrix of model coefficients at lag *k* whose *I,j* element is *a*_*i,j*_*(k)*, and **u** (*n*) is the *M-by-1* vector of model errors or innovations. We shall assume that Eq. (13) represents the rPCA case as well, with the appropriate substitutions of the virtual electrode data.

#### Mean-square Prediction Error

Let z*(n)* denote new simulated data and **Â**(*K*) the estimated MVAR model coefficients. The mean square prediction error is

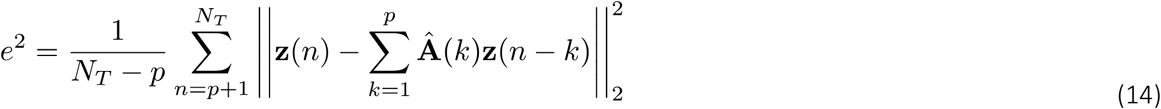

where *NT* is the number of simulated test samples corresponding to 16 minutes of data.

#### Partially Directed Coherence Measures

Many different measures of connectivity have been proposed for MVAR models. Here, we used a variant of the generalized partial directed coherence (gPDC) to characterize the connectivity between ROIs, i.e., vector time series, with a scalar metric. Eq. (13) may be expressed in the frequency domain as

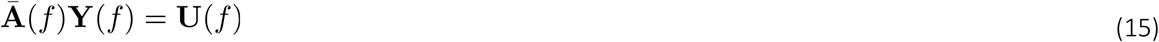

where

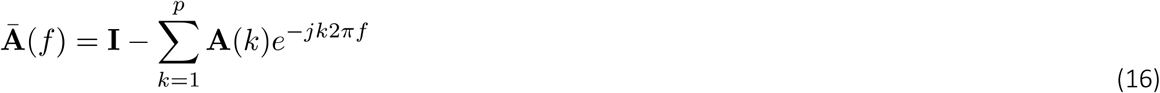

Here 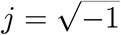 and *f*is frequency normalized by the sampling frequency, i.e., -0.5 < *f*≤ 0.5 with units of cycles per sample. Further, let superscript *H*denote the complex conjugate transpose operator, **Σ = *E* {*U*(*f*) *U***^***H***^ **(*f*)}** denote the error covariance matrix, assumed constant over frequency, and **Φ = Σ**^**−1**^ the inverse error covariance matrix. The gPDC metric (Baccala et al., 2007) from electrode *j* to electrode *i* is written

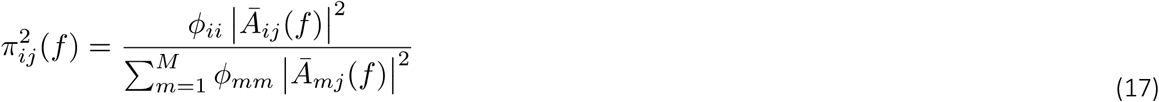

where *ϕ*_*kk*_ is the *k*^*th*^ diagonal element of **Φ** and **Ā**_*ij*_ (*f*) is the *I,j* element of **Ā**. (*f*) Similarly, we define the block gPDC for the relationship from the *j*^*th*^ vector time series to the *i*^*th*^ vector time series as

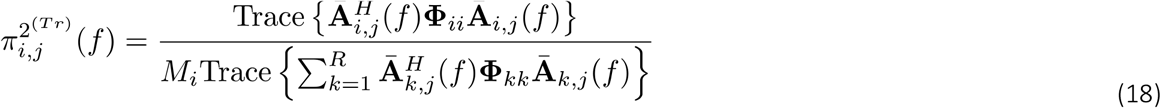

where **Ā**_*ij*_ (*f*) is the block of **Ā**. (*f*) associated with the connection from the *j*^*th*^ to *i*^*th*^ vector time series, **Φ**_*ii*_ is the block of **Φ** associated with the *i*^*th*^ vector time series, and the matrix trace operation is the sum of the diagonal elements. Note that the factor *M*_*i*_ normalizes the block gPDC metric by the number of electrodes associated with the target ROI. The relationship between the measure in Eq. (18) and the previously proposed vector connectivity measures of (Faes and Nollo, 2013) is described in Supplementary Methods.

We computed broad band connectivity by integrating 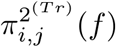 over frequency. Define the ground truth broad band connectivity from electrode *j* to *i* as *c*_*i,j*_ and the estimated broad band connectivity in trial *k* as 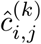. The acronym bgPDC (block gPDC) is used to refer to broadband connectivity *c*_*i,j*_ and 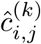. We compared the ground truth bgPDC to the mean estimated bgPDC over ten trials visually (e.g., see Figure 4, Supplementary Figures 3-6). As a global quantitative measure of fidelity, for each subject and data length we computed the mean-absolute-difference connectivity between the ground truth and estimated bgPDC connectivity, defined as

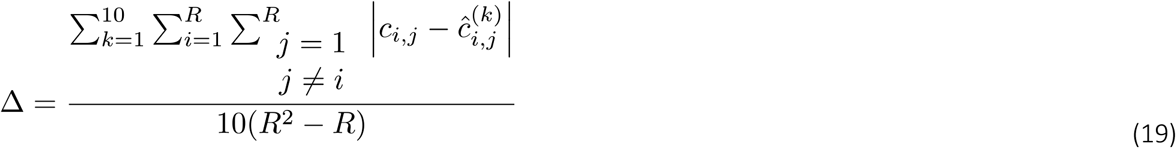

## Results

### Effective connectivity in ground truth networks

Resting state iEEG recordings were obtained in ten neurosurgical patients (Figure 1A; Supplementary Figure 1; Table 2). MVAR model fits to 4-minute data segments were used to derive effective connectivity profiles (Figure 1C-D). Connectivity matrices at the single electrode and ROI levels displayed strong symmetry. In the connectivity matrices of Figure 1, the electrode locations and ROIs are ordered in a roughly hierarchical fashion, so as expected, connectivity was strongest along the diagonal.

To explore further the consistency of these results with previous reports, we evaluated the connectivity patterns in all subjects to and from the anterior and posterior subdivisions of the insula (Figure 3), areas that are anatomically close to each other with distinct functions but strong interconnectivity (Augustine, 1996; Cauda et al., 2011; Cloutman et al., 2012; Zhang et al., 2019). The results of Figure 3 are consistent with these previous reports. InsP is considered a sensory region, both interoceptive and exteroceptive, and in the ROIs sampled would be expected to have strong connections to early auditory cortical structures. This is evident in the analyzed dataset as strong bright yellow vertical bands in Figure 3A corresponding to ROIs (HGPM, HGAL) on Heschl’s gyrus, the location of primary auditory cortex. By contrast, InsA is a higher order structure with strong connectivity to the higher order auditory cortical structure planum polare (PP; Figure 3B). Both InsP and InsA sent strong projections to amygdala. Additionally, bidirectional connectivity was observed between InsP and InsA, with a higher connection strength from InsA to InsP.

**Figure 3.**
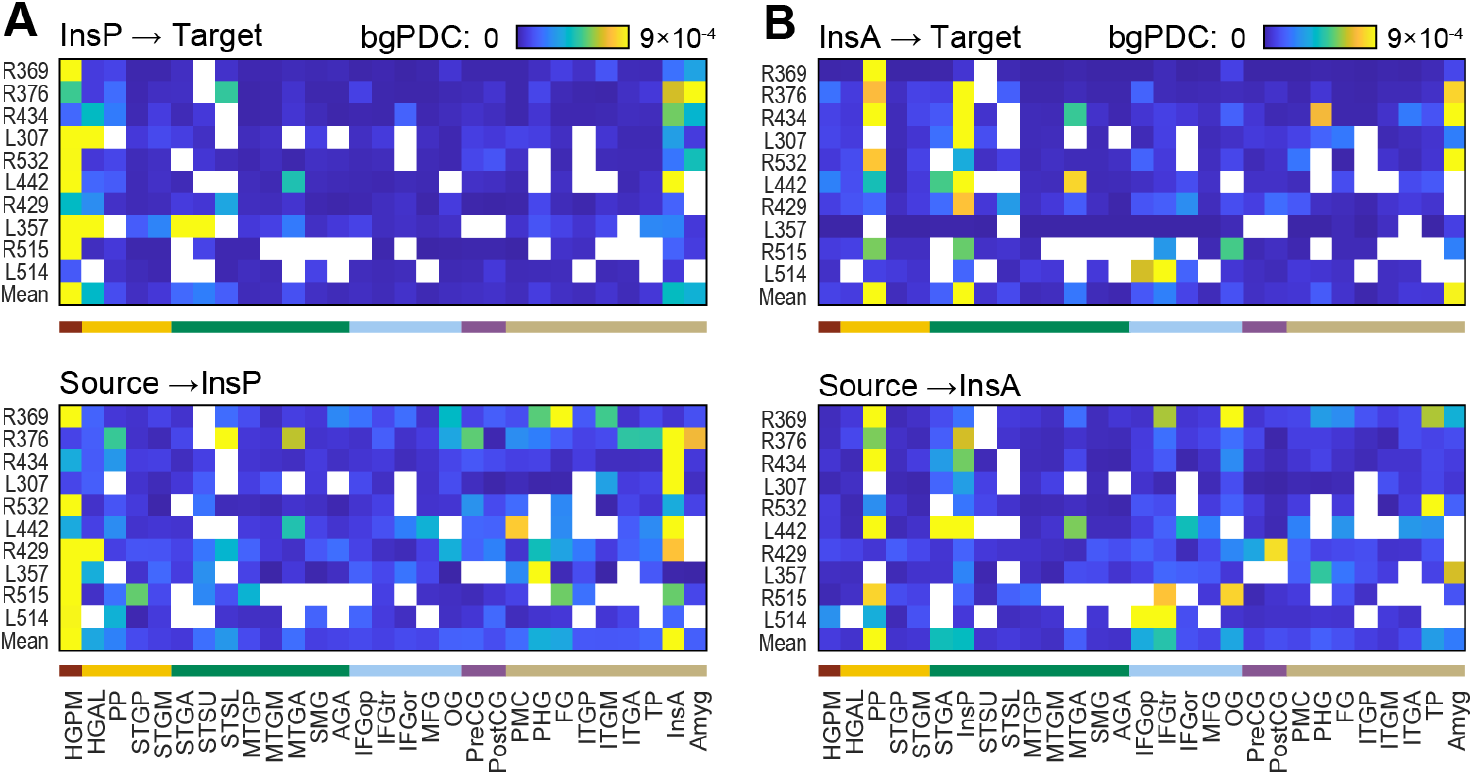
Summary of connectivity of insula for all subjects. A: Posterior insula (InsP). B: Anterior insula (InsA). Only ROIs with coverage in 6 or more subjects are shown. Color bars underneath each panel represent ROI groups, color-coded as shown in the legend of Figure 1A. See Table 2 for list of ROI abbreviations.

### Simulation Experiment

The performance of the four MVAR model estimation methods was evaluated using model fits to data of varying lengths simulated from ground truth models (Figure 4). We report the data length *T* in s to simplify the application of the results to the original subject data; with the sampling rate fixed at 250 s^-1^, the number of samples *N = 250T* In the example shown, the OLS and rPCA estimates failed to capture any features of the ground truth connectivity matrix for *T* = 10 s, in contrast to rPCA-gLASSO and gLASSO. Modest improvement was observed for PCA at *T* = 30 s. When *T* was increased to 60 s, the performance of the OLS and rPCA estimates improved, but connectivity tended to be substantially overestimated for both methods (Figure 3). Comparable results were observed in the ground truth models from all subjects (Supplementary Figures 3-6). This is illustrated more directly in Figure 5, which shows scatterplots of estimated vs. true connectivity for a single subject for 10, 30, and 60 s. Only the strongest connections were estimated correctly with OLS and rPCA estimates even for *T* = 60 s. There was a slight overestimation of connectivity values for rPCA-gLASSO and gLASSO estimates as well, but in general these estimates were much more accurate. An example from a second subject is shown in Supplementary Figure 7. Model performance was quantified across subjects using mean-square prediction error (Eq. 14) and bgPDC fidelity c_*MAD*_ (Eq. 19) as shown in Figure 6. As *T* increased, both metrics improved for all estimators. However, the rPCA-gLASSO and gLASSO estimates performed substantially better than the OLS estimate in the most data-limited case (*T =* 10 s). In fact, a data length of *T* = 960 s was required for OLS to show superior one-step prediction error compared to gLASSO (Figure 6B). We note as well that while the rPCA-gLASSO and gLASSO estimates performed comparably to each other on these metrics, the gLASSO model required significantly more computation to estimate.

**Figure 4.**
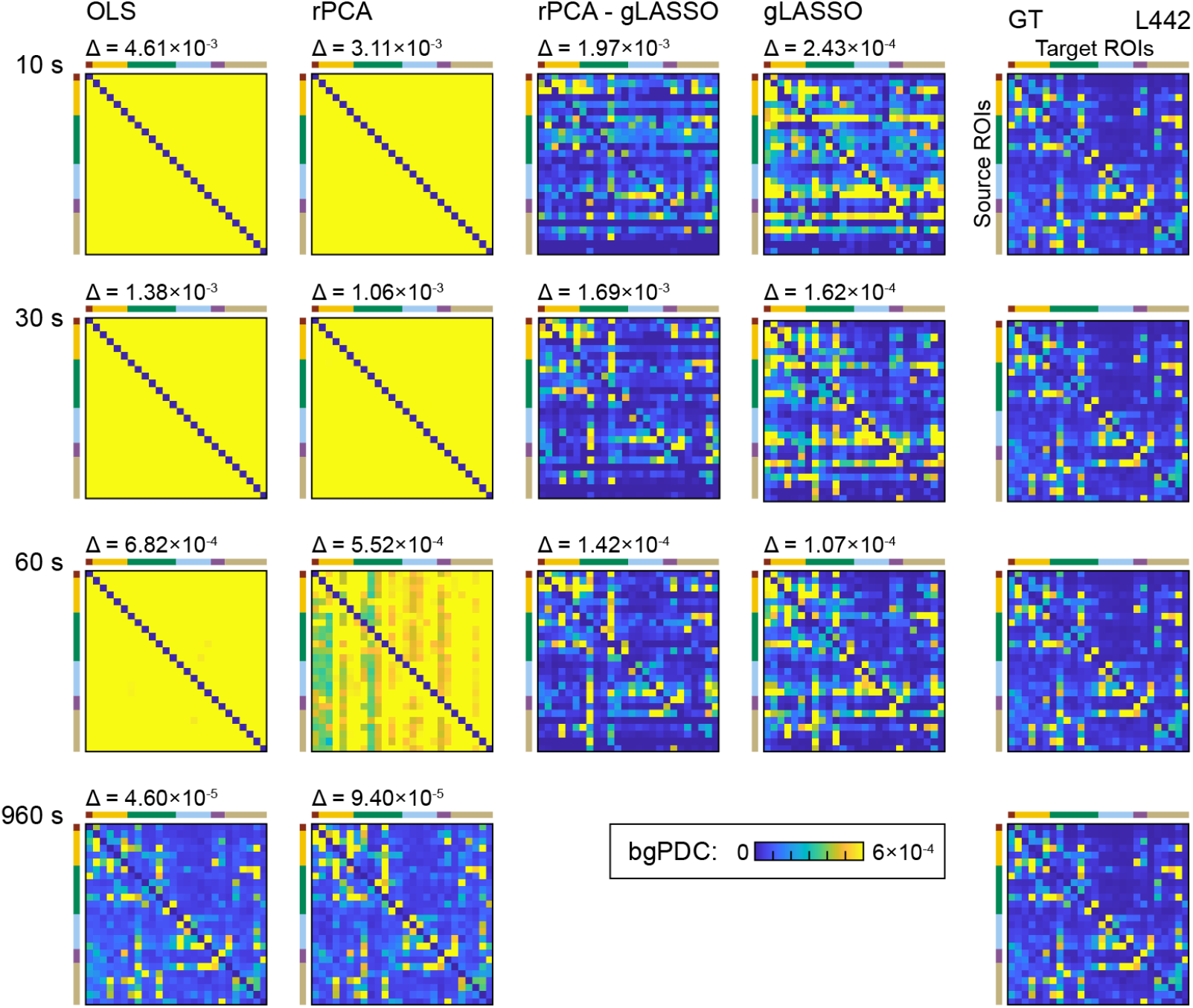
Recovery of ground-truth ROI connectivity (bgPDC) in subject 442L. Adjacency matrices show rows as origin ROIs, and columns as target ROIs. Each connectivity matrix depicts the average result over ten trials, except for the *T = 960* s case, which represents an average of two. The delta values displayed above each recovery matrix represent the mean absolute error of the average recovery matrix compared to the ground-truth (GT) matrix as defined in Eq. (19). The same color scale is used for all adjacency matrices shown. The maximum color value is based off of the 95^th^ percentile of the ground-truth matrix. Color bars next to the connectivity matrices represent ROI groups, color-coded as shown in the legend of Figure 1A.

**Figure 5.**
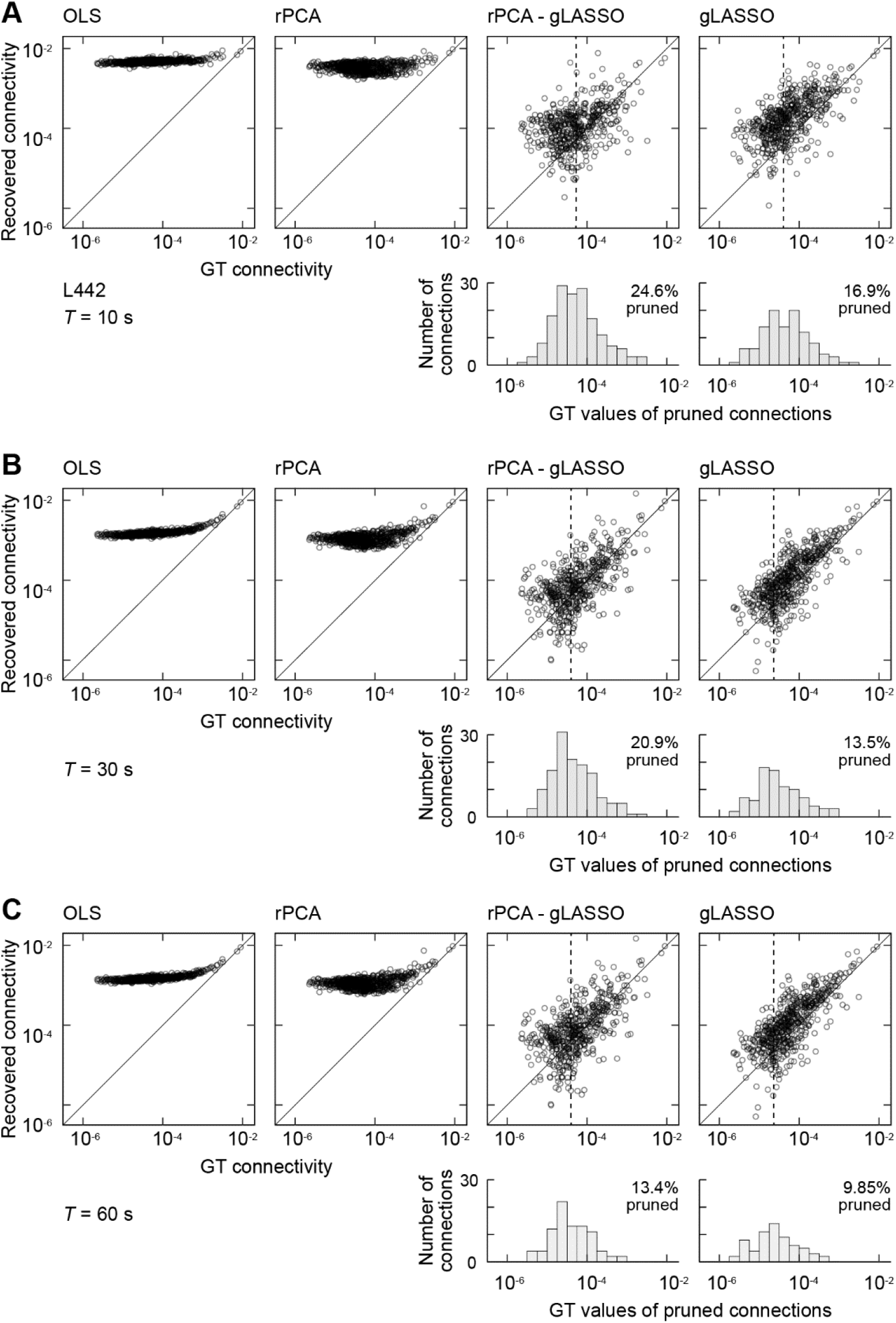
Comparison of ground-truth vs recovered bgPDC connectivity. Scatter plots of average (10 trials) estimated versus ground truth connectivity for subject 442L for different estimation methods. The rPCA-gLASSO and gLASSO estimation results include histogram of connections that are pruned to zero by the gLASSO procedure. The dashed vertical line in the histograms denotes the median of the ground truth connectivity values that have been pruned. A Connectivity estimated for *T* = 10 s of simulated data. **B** *T = 3*0 s. **C**. *T* = 60 s.

**Figure 6.**
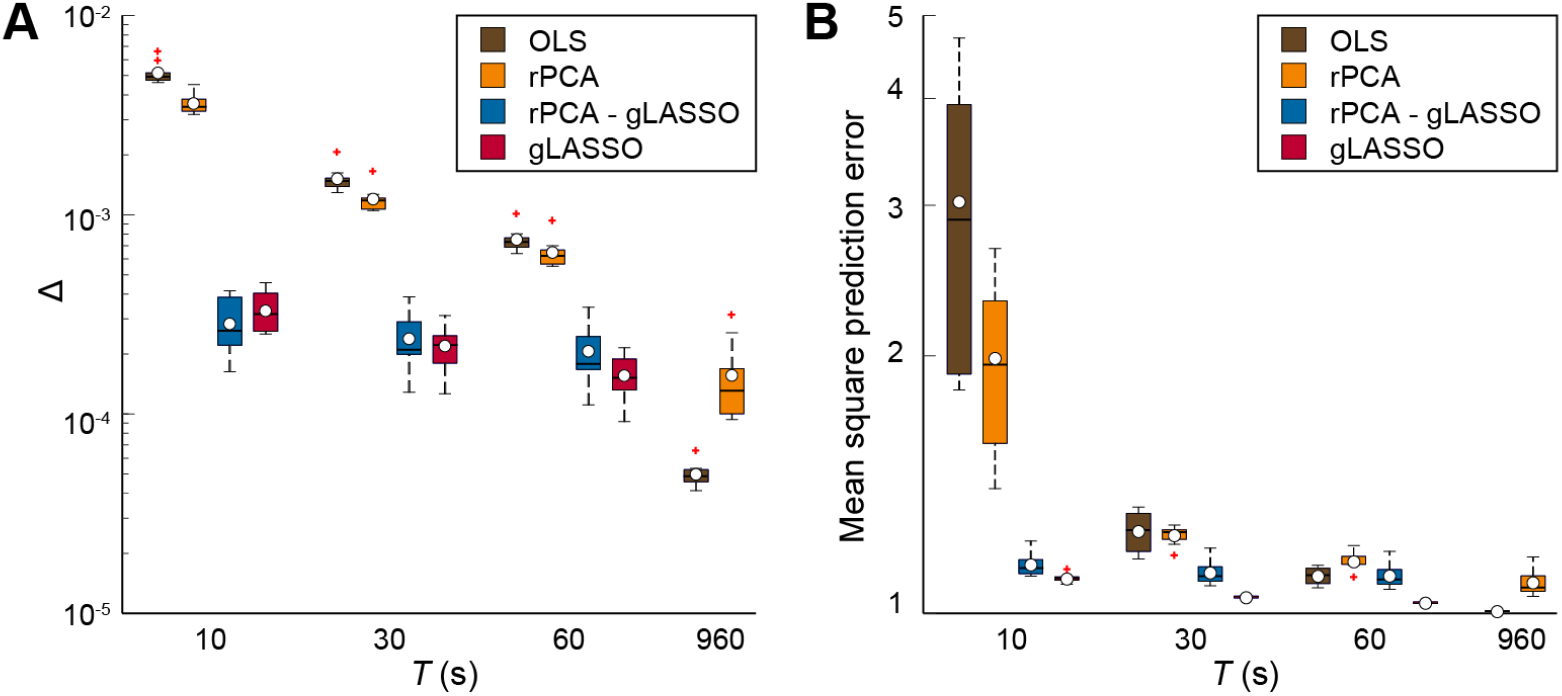
Model performance. Summary of estimation performance as a function of data length across all simulated ground truth models. The results for each model are averaged over ten trials and the box plots characterize average performance across the ten models. A. Mean-absolute-difference connectivity between the ground truth and estimated bgPDC connectivity (Δ; Eq. 19). B. One-step mean-square prediction error (Eq. 14). Box and whiskers plots depict medians (horizontal lines), means (white symbols), quartiles (boxes), ranges (whiskers) and outliers (red crosses).

**Figure 7.**
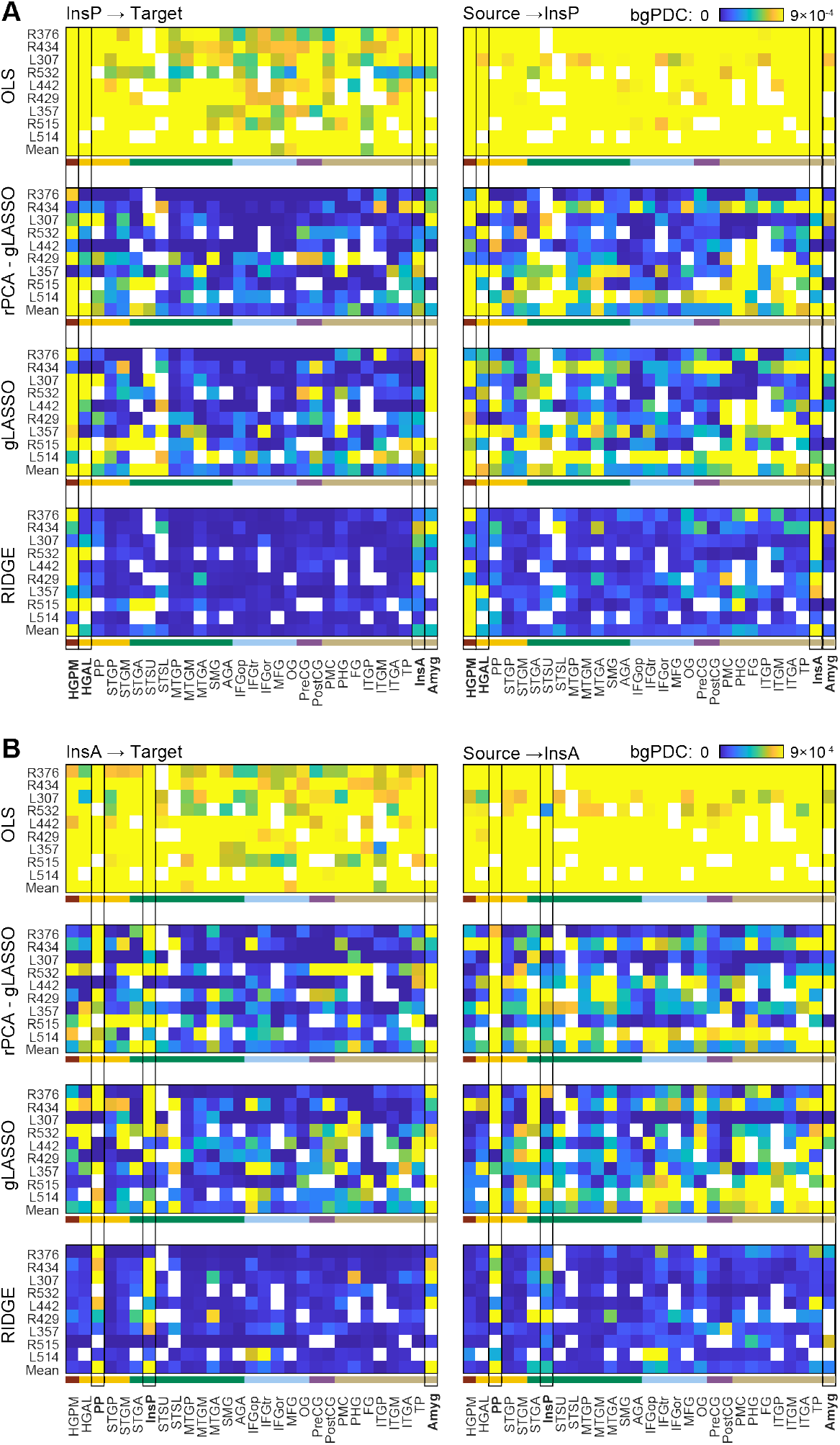
Connectivity estimated from human subject data. Estimated bgPDC connectivity from (*left*) and to (*right)* posterior insula (A) and anterior insula (B). Each column subplot represents a different modeling strategy applied to recorded data that was used to estimate the original 10 GT models. The RIDGE models were fit to four minutes of resting-state data. To determine if we can recover these models using limited data, we applied each modeling approach to four one-minute segments of the human subject data. Results are averaged across all four segments for each subject/model-type. Highlighted ROIs are discussed specifically in the text. Color bars underneath each panel represent ROI groups, color-coded as shown in the legend of Figure 1A. See Table 2 for list of ROI abbreviations.

### Plausibility Analysis

The data of Figures 4 – 6 indicate that gLASSO and rPCA-gLASSO estimates provided superior performance in capturing known connectivity from ground-truth models. We next explored the performance of OLS, gLASSO, and rPCA-gLASSO estimation methods when applied to one-minute segments of the original human subject data that was used to generate the ground-truth models. Each estimation method was applied to four 1-minute segments of the human subject data. The ground truth connectivity was not known, so in Figure 7 we compare bgPDC connectivity profiles to and from the anterior and posterior subdivisions of the insula in all subjects to assess consistency with the results for longer data segments shown in Figure 1 and with previous reports from the literature. The severe overestimation of connectivity associated with the OLS method was clearly evident, as was the more modest overestimation with gLASSO and rPCA-gLASSO. In general, the latter two approaches captured the differential connectivity of InsP and InsA to early (HGPM, HGAL) versus higher order (PP) auditory structures and the amygdala, indicating that results were plausible even in the absence of ground truth knowledge.

### Dynamics of iEEG connectivity

The data of Figure 5 and Supplementary Figure 7 indicate that gLASSO and rPCA-gLASSO can reliably estimate connectivity even for short data lengths (= 10 s). The possibility of estimating high-dimensional network models from relatively short data segments enables the study of changes in connectivity that occur over time, a growing area of interest in research on brain networks (Preti et al., 2017). To illustrate the importance of performing dynamic connectivity analysis with high temporal resolution, we explored dynamic connectivity during ten minutes of resting state iEEG data in two subjects using the gLASSO estimation method with models based on = 60 and 10 s (Figure 8). To visualize changes in connectivity over time, we stacked the connectivity estimates between all ROIs into a single vector and sorted the connectivity from smallest to largest mean connectivity over the ten minutes. Sorted connectivity is displayed on the vertical axis and time on the horizontal axis. Thus, each row depicts the temporal evolution of connectivity between two ROIs. To further identify potential temporal patterns, we applied singular value decomposition to this matrix of vectorized connectivity vs time and extracted the dominant component. The right-most figure in each panel shows the dominant component with the time course of the dominant component depicted beneath. The results based on a 10-s segment revealed temporal patterns that were smoothed over by the longer 60-s window. In some cases, rapid changes in connectivity were completely obscured for the 60-s models. Thus, methods to facilitate accurate representations of network connectivity over short times scales are essential to capture the dynamics of the brain in resting state.

**Figure 8.**
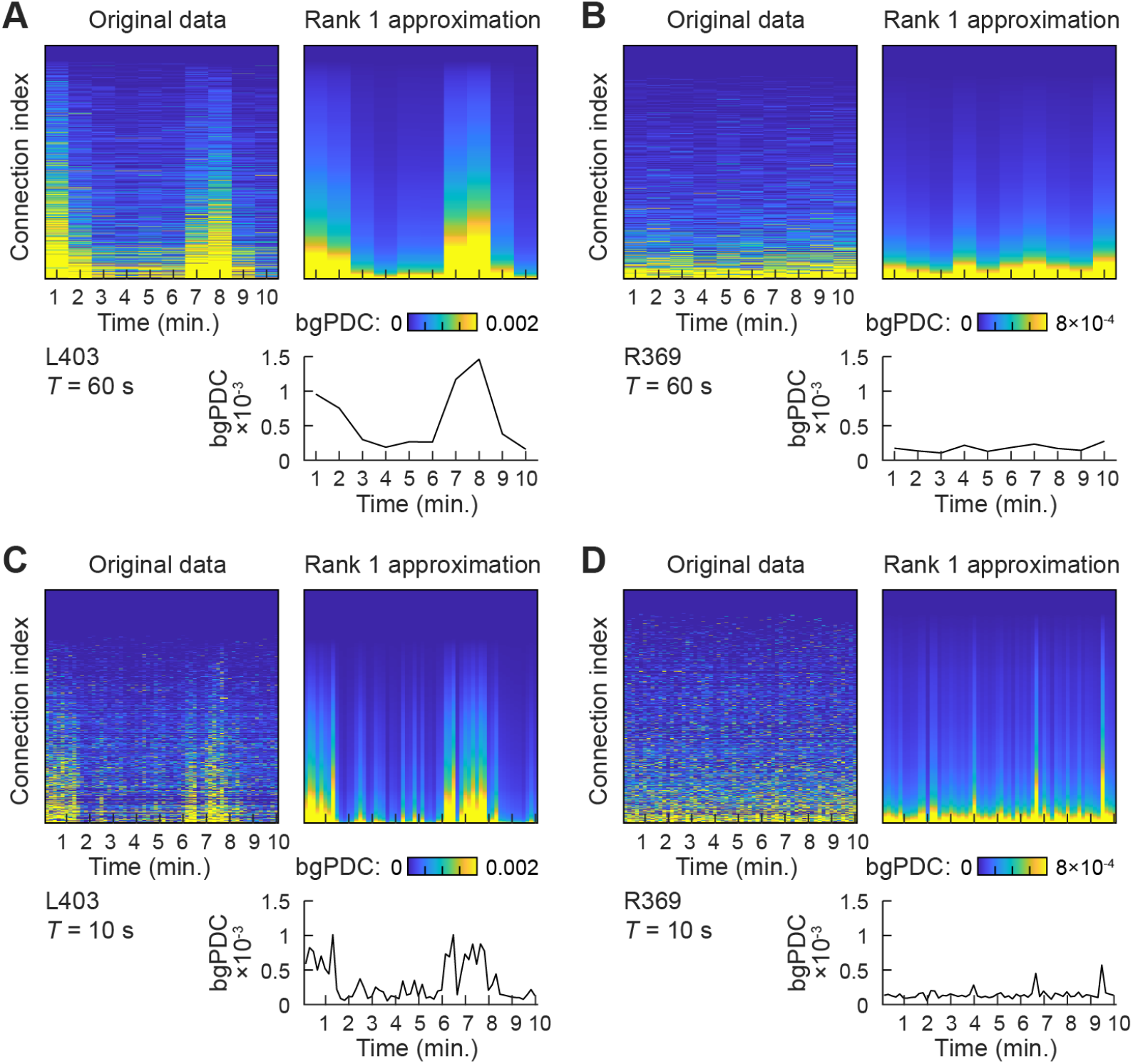
Connectivity dynamics. Each panel depicts the gLASSO-estimated sorted dynamic connectivity matrix on the left and the best rank-1 approximation on the right. In each panel, the line plot below the rank-1 approximation depicts the mean across connectivity indices to illustrate the overall temporal evolution. The estimates are derived from *T* = 60 s (A, B) and *T* = 10 s (C, D) segments of human subject data. A,C: Subject L403. B,D: Subject R369.

## Discussion

It has been widely noted in the literature that long recording durations are needed to estimate high-dimensional MVAR models in time series data. For example, Schlogl and Supp (Schlogl and Supp, 2006) propose that the number of data samples should exceed ten times the product of the memory times the number of electrodes (10*Mp*). Such requirements likely explain why MVAR models have not been reported in the literature for more than ∼30 electrodes. Our human subject and experimental results with OLS model estimation indicate that the factor of ten suggested may be too small for networks in the 100-200 electrode range. We found that four minutes (∽60,000 samples) of human subject based resting-state data proved inadequate for OLS estimation of MVAR models with ∽100-200 electrodes and memory of eight (*Mp* ∼ 1600), which is why we employed ridge regression to estimate ground-truth models from four minutes of data. Our simulation studies show that the OLS approach consistently overestimates connectivity for *T* = 60 s (Figures 4 and 5; Supplementary Figures 3-6), corresponding to a data length of 15,000 samples. For this dataset, 10*Mp* ranged from 9360 to 17,280 (average = 13,520. Clearly there is a need for estimation methods that are effective with less data.

The overestimation of connectivity observed with the OLS approach is indicative that the matrix inverse required to compute Eq. (4) is ill-conditioned. The *Mp*-by*Mp*-matrix **Y**^***T***^**Y** is an estimate of the covariance matrix of the data. Few statistical results are available for the properties of this matrix due to the temporal dependence of the columns of **Y**. However, if the columns of **Y** are independent – a more stringent condition – and multivariate Gaussian, then **Y**^***T***^**Y** is a sample covariance matrix with a Wishart distribution (Muirhead, 1982). Such matrices have been studied extensively and the distributions of the eigenvalues are known (Muirhead, 1982), though they are complex and challenging to interpret. Asymptotic distributions of eigenvalues follow the Marchenko-Pastur law (Pastur and Marchenko, 1967). For our purposes, it is sufficient to note that the small eigenvalues of a sample covariance matrix require much more data for reliable estimation than the large eigenvalues. Inversion of **Y**^***T***^**Y** involves inverting the eigenvalues, so the small, underestimated eigenvalues of **Y**^***T***^**Y** become dominant in, resulting in potentially very large noisy contributions to the OLS weights in Eq. (4). This effect is mitigated by ridge regression, which is why we chose ridge regression for estimating ground truth models. Addition of γ^**I**^ in Eq. (6) ensures all eigenvalues of the matrix being inverted exceed γ, thus limiting the magnification of noisy contributions in the model weights. The downside of using ridge regression is that the model squared error increases as γ increases.

One approach to enabling MVAR modeling of large networks is to reduce the inherent dimensionality of the problem. We applied PCA to regional collections of electrodes to create a smaller number virtual electrodes that represent the regional activity. While the rPCA approach modifies interelectrode connectivity, it preserves regional connectivity provided relevant activity is represented in the principal components. Although rPCA maximizes the variance represented in the region for a given dimension, interregional connectivity bias is not necessarily proportional to variance represented. It is theoretically possible that a smaller PCA component discarded in the rPCA process makes a significant contribution to interregional connectivity. This potential is illustrated by the 960-s results shown in Figure 6. The performance of rPCA is noticeably worse than the OLS estimates, reflecting the bias incurred by discarding five percent of the variance. The rPCA approach cannot predict components that have been discarded. However, Figures 4-6 indicate that the rPCA approach is less sensitive to overestimation artifacts than the OLS method with shorter data record lengths. Hence, there is a tradeoff between the connectivity bias incurred by discarding dimensions and the reduction in estimator variance associated with smaller problem size. Concerns over connectivity bias led us to explore retention of 95 percent of the variance in each region, which limited the dimensionality reduction achieved via rPCA to a factor of about two in our data.

The gLASSO regularizer is motivated by the expectation of sparse connectivity in the brain, such as that associated with small world networks (Achard et al., 2006; Bullmore and Sporns, 2009; Sporns and Zwi, 2004), and effectively reduces the dimensionality of the problem by “turning off” or pruning connections that do not have a significant impact on the squared error for a given data length. The potential connectivity bias associated with pruning connections is justified because these connections have the weakest impact on the squared error for the available data as determined by cross validation. This procedure ensures that the reduction in estimator variability due to reduced dimensionality outweighs potential bias for the data set. The histograms in Figure 5 indicate that weaker connections are most likely to be turned off by gLASSO. Note that the gLASSO approach results in sparser models as the data length decreases (see, e.g., Figures 4 and 5) because it becomes more difficult to establish the significance or presence of additional connections as the data length available for training the model decreases. We note, however, that these benefits of the gLASSO method accrue at the expense of significant computational cost.

The rPCA-gLASSO approach has a lower computational burden than the gLASSO method due to the reduction in dimensionality associated with rPCA. Since the model coefficients associated with predicting any one electrode can be estimated as an individual optimization problem, reducing the effective number of electrodes by a factor of two will cut the total computational runtime of solving an MVAR model in half—independent of the method used to solve for each electrode’s associated coefficients. This improvement becomes considerably more noticeable when hyperparameter tuning methods are incorporated into the model-fitting step, such as cross-validation, to optimize the sparsity hyperparameter (λ) for each electrode in a gLASSO model. However, the rPCA-gLASSO method is subject to the connectivity bias discussed in the rPCA approach. Furthermore, rPCA methods only apply to regional connectivity estimation involving multiple electrodes sampling a region. If electrode-by-electrode connectivity is needed, then rPCA is not applicable.

Our simulation study enables objective evaluation of estimator performance by comparison to the ground truth for a biologically realistic model. We chose to quantify performance using mean-square prediction error (Eq. 14), mean absolute deviation of bgPDC connectivity (Eq. 19), and estimated versus ground truth bgPDC (Figure 5). There are many different metrics that could be used to measure performance as well, and different metrics may show different sensitivity to estimation errors. For example, the mean absolute deviation of bgPDC connectivity is more sensitive to data length than the one-step mean-square prediction error. Overestimation of connectivity affects bgPDC more than mean-square prediction error on the test data, which is to be expected since the OLS procedure minimizes the prediction error on the training data. It is likely that multiple connectivity patterns could produce comparable prediction error, especially with relatively short data records. The OLS estimator is biased toward overestimation of connectivity. In effect it produces large connectivity values that tend to offset one another and give reasonable squared prediction error.

We chose to consider only a fixed model order of *p = 8*. This simplified comparison of results for ground truth models associated with different subjects and reduces potential difficulties of interpreting performance across ground-truth models with different model order. Model order has a direct impact on model complexity and model-order selection can be viewed as a form of regularization. Low data scenarios call for lower model orders than those supported by higher data scenarios. This dependence is reflected in commonly used order-selection strategies such as the Akaike Information Criterion (Cavanaugh and Neath, 2019) and in the recommendation that the data length be >10*Mp* (Schlogl and Supp, 2006). By fixing the order of our estimated models, we can isolate the impact of the estimation method on estimation quality.

We implemented rPCA to reduce dimensionality in the data but still retain spatial information. Alternatively, PCA can be run on the full time-series dataset followed by projecting the PCA-based model residuals back to a full-dimensional representation (Schmidt et al., 2016). The disadvantage of the latter approach is that it precludes considering the regional structure, which is needed to assess regional connectivity. Additionally, the benefit of the rPCA method over the projection method arises due to the inability to project the model’s coefficients back to a high-dimensional state—meaning that only the error terms of the projected model can be used for estimation of connectivity. Since the calculation of the very popular connectivity metric—the gPDC—requires the use of model coefficients, the rPCA method is advantageous in this domain of work.

Application of these estimation methods to human subject data suggests that plausible connectivity estimates are obtained with *T* = 60 s and *M* = 200 electrodes (Figure 7). We chose to explore connectivity to and from anterior versus posterior insula, due to this region’s importance in intero- and exteroceptive sensory processing, salience, emotion, homeostasis, and consciousness (Huang et al., 2021; Zhang et al., 2019). Although anatomically adjacent, anterior and posterior insula have distinct functional roles and connectivity profiles (Cauda et al., 2011). For example, the anterior portion of the insula is thought to couple tightly with prefrontal regions and amygdala. The posterior portion, on the other hand, is more closely linked to activity propagated from early auditory regions—including Heschl’s gyrus (Zhang et al., 2019). The results of Figure 3 are consistent with these functional distinctions. The strong connections identified in the present analysis are also consistent with results of tracer injection studies (reviewed in (Augustine, 1996)). In addition, the relatively strong connectivity of InsA with the IFG and rostral superior temporal regions STGA and PP is consistent with previous probabilistic tractography results (Cloutman et al., 2012). Effective connectivity derived from MVAR model fits was consistent with these previous studies. Even for shorter data lengths, model fits relying on gLASSO were able to capture expected connectivity profiles and community structure in brain networks, for example demonstrating strong connectivity between auditory cortical structures (upper left quadrants in the hierarchically-sorted adjacency matrices shown in Figure 4 and Supplementary Figures 3-6). Motivated by the reasonable recovery of GT connectivity for *T* = 10 s demonstrated in Figure 5 and Supplementary Figure 7, we also show that models fit to short data lengths (*T* = 10 s) can capture rapid dynamics of network connectivity that are obscured at a resolution of *T* = 60 s. Importantly, given the likelihood that brain activity is nonstationary except over very brief intervals, the capability to fit models to short data segments allows for more accurate piecewise-stationary estimates of dynamic brain activity.

The results indicate that the rPCA-gLASSO and gLASSO methods reliably estimate MVAR models for large networks with far less data than previously thought possible. The capability to estimate models for networks with large numbers of electrodes reduces the likelihood of detecting spurious effective connections resulting from removed or missing mediating electrode channels and leads to improved connectivity analyses. Furthermore, reducing data requirements reduces concerns about nonstationarity in the data, and creates new analysis opportunities, such as assessment of dynamic connectivity. Consequently, the methods presented here will have broad application and substantial impact for interpreting resting state data recorded from human subjects, as well as large scale multielectrode recordings in experimental animals.

## Acknowledgements

This research was performed using the computer resources and assistance of the UW-Madison Center for High Throughput Computing (CHTC) in the Department of Computer Sciences. The CHTC is supported by UW-Madison, the Advanced Computing Initiative, the Wisconsin Alumni Research Foundation, the Wisconsin Institutes for Discovery, and the National Science Foundation, and is an active member of the OSG Consortium, which is supported by the National Science Foundation and the U.S. Department of Energy’s Office of Science. This research was done using resources provided by the Open Science Grid (Pordes et al., 2007), which is supported by the National Science Foundation award #2030508. Funding provided by NIH/NIGMS (R01 GM109086 to MIB & KVN) and the UW Department of Anesthesiology (to MIB).

## Data and code availability

Data is available via a request to the Authors pending establishment of a formal data sharing agreement, submission of a formal project outline, and agreement about co-authorship. All software developed for this study will be freely available via Git repository. Please contact Bryan Krause for details.

