## Supplementary figures and appendix for "Multivariate autoregressive model estimation for high dimensional intracranial electrophysiological data"

Author contributions: Christopher Endemann: Conceptualization, Methodology, Software, Formal analysis, Investigation, Writing - Original Draft, Visualization; Bryan M. Krause: Conceptualization, Software, Data Curation, Writing - Original Draft; Kirill V. Nourski: Investigation, Data Curation, Writing - Review & Editing, Visualization, Funding acquisition; Matthew I. Banks: Conceptualization, Investigation, Writing - Original Draft, Supervision, Project administration, Funding acquisition; Barry Van Veen: Conceptualization, Methodology, Writing - Original Draft, Supervision, Project administration.

Acknowledgements: This research was performed using the computer resources and assistance of the UW-Madison Center for High Throughput Computing (CHTC) in the Department of Computer Sciences. The CHTC is supported by UW-Madison, the Advanced Computing Initiative, the Wisconsin Alumni Research Foundation, the Wisconsin Institutes for Discovery, and the National Science Foundation, and is an active member of the OSG Consortium, which is supported by the National Science Foundation and the U.S. Department of Energy's Office of Science. This research was done using resources provided by the Open Science Grid (Pordes et al., 2007), which is supported by the National Science Foundation award #2030508. Funding provided by NIH/NIGMS (R01 GM109086 to MIB & KVN) and the UW Department of Anesthesiology (to MIB).

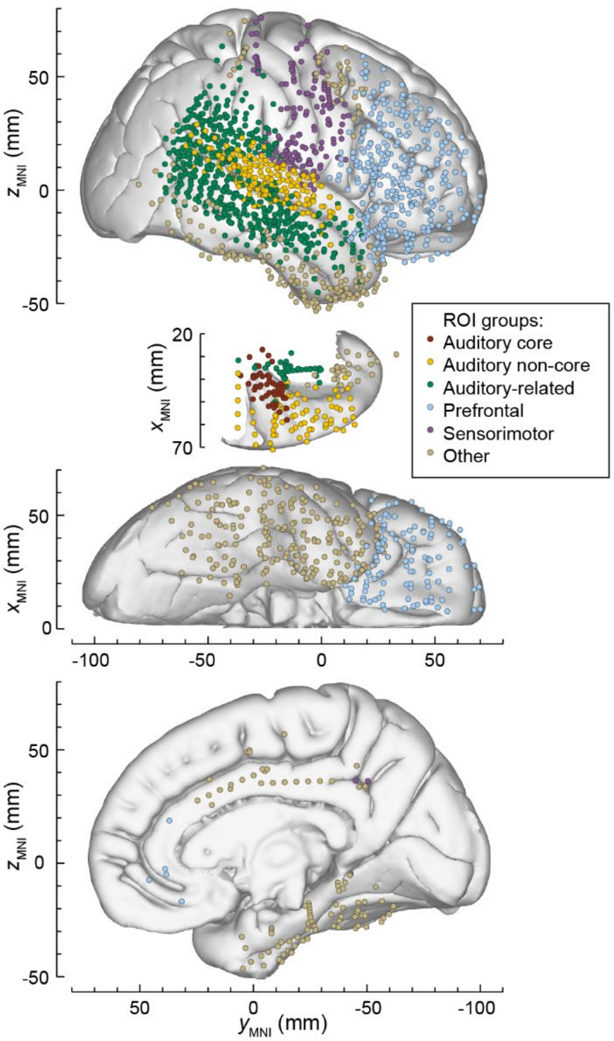


**Supplementary Figure 1.** Electrode coverage in all subjects. Locations of recording sites, color-coded according to the ROI group, are plotted in MNI coordinate space and projected onto the FreeSurfer average template brain for spatial reference. Left hemisphere MNI *x*-axis coordinates ($x_{MNI}$) of recording sites in subjects L307, L357, L442, and L514 were multiplied by -1 to map them onto the right-hemisphere common space. Projection is shown on the lateral, top-down (STP), ventral and mesial views (top to bottom rows). Recording sites over orbital, transverse frontopolar, inferior temporal gyrus and temporal pole are shown in both the lateral and the ventral view. Sites in fusiform, lingual, parahippocampal gyrus and gyrus rectus are shown in both the ventral and medial view. Sites in the amygdala ($n$ = 19), caudate nucleus ($n$ = 1), frontal operculum ($n$ = 6), hippocampus ($n$ = 16), parietal operculum ($n$ = 4), putamen ($n$ = 6), substantia innominata ($n$ = 2), and ventral striatum ($n$ = 2) are not shown.

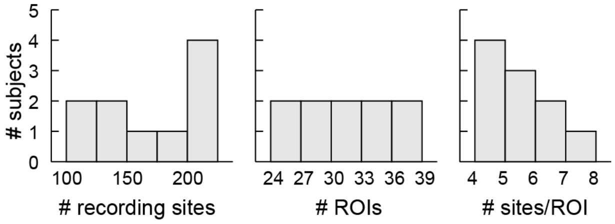


Supplementary Figure 2. Histogram showing descriptive statistics for the 10 subjects used in the simulation experiment.

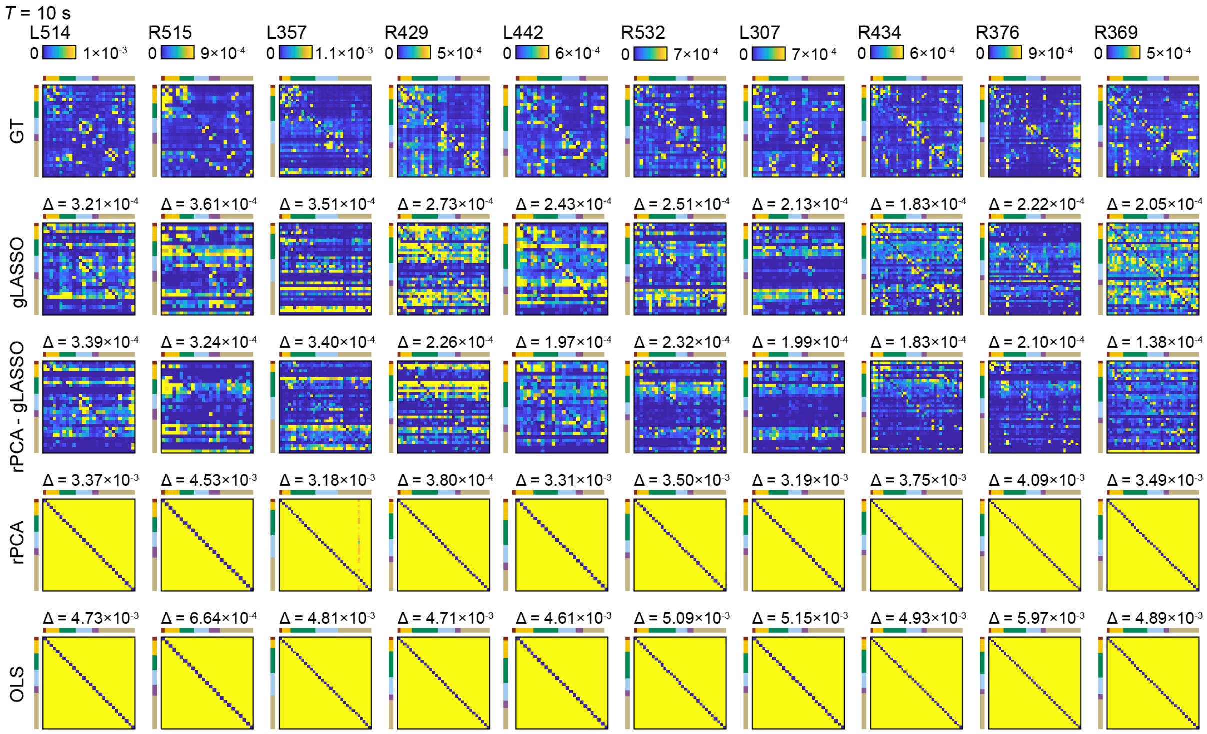


**Supplementary Figure 3.** Recovery of ground-truth ROI connectivity (bgPDC) in all subjects for $T$ = 10 s. Adjacency matrices show rows as origin ROIs, and columns as target ROIs. Each connectivity matrix depicts the average result over ten trials. The delta values displayed above each recovery matrix represent the mean absolute error of the average recovery matrix compared to the ground-truth (GT) matrix as defined in Eq. (19). The same color scale is used for all adjacency matrices shown. The maximum color value is based off of the 95th percentile of the ground-truth matrix. Color bars next to the connectivity matrices represent ROI groups, color-coded as shown in the legend of Figure 1A.

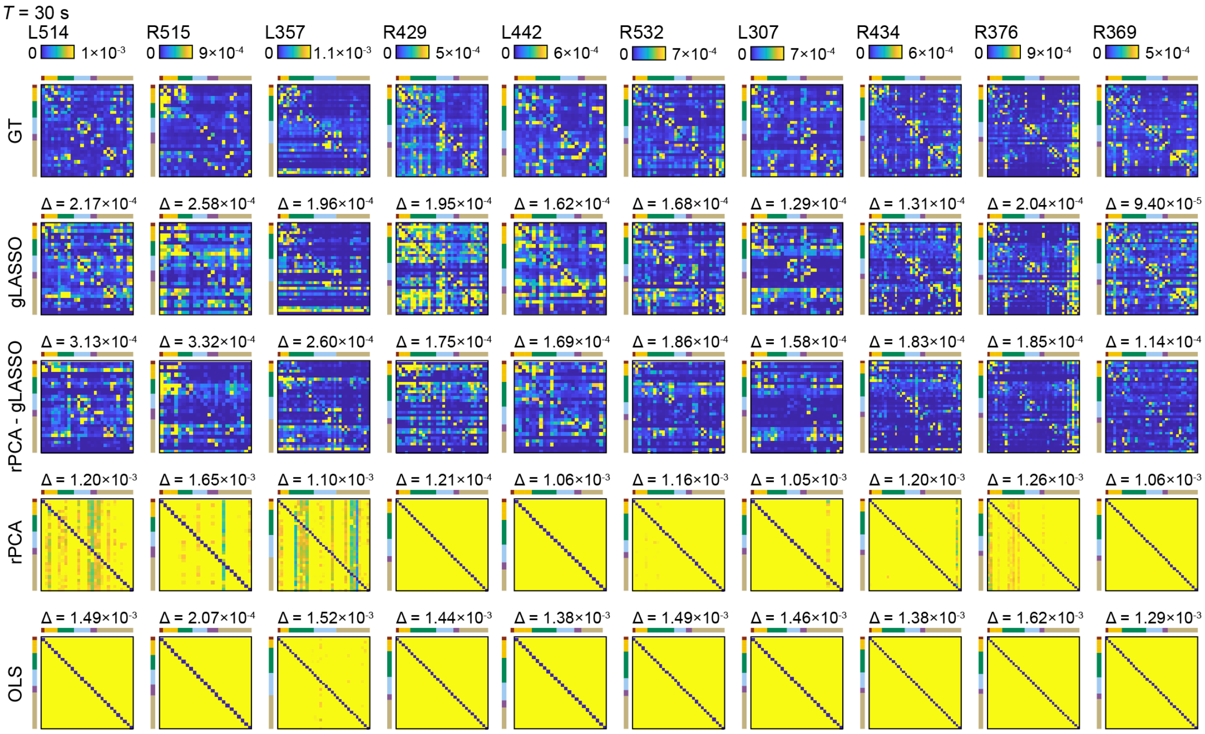


**Supplementary Figure 4.** Recovery of ground-truth ROI connectivity (bgPDC) in all subjects for $T$ = 30 s. See caption of Supplementary Figure 3 for detail.

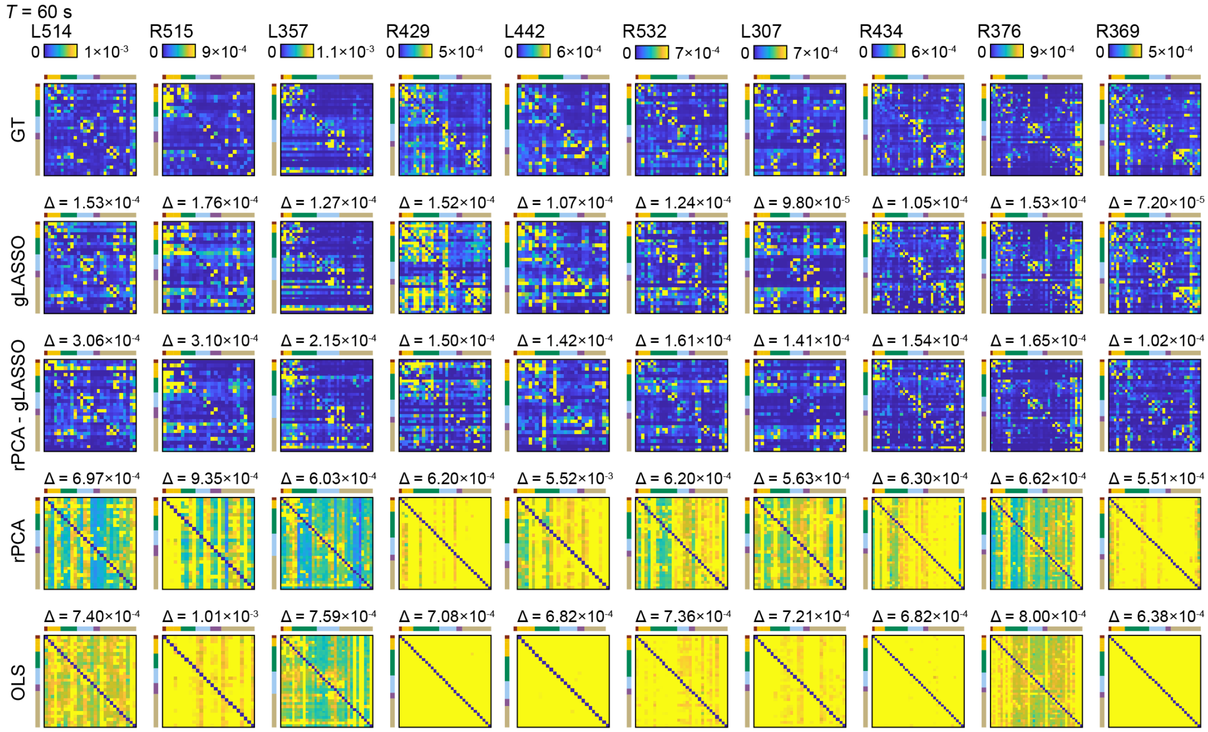


**Supplementary Figure 5.** Recovery of ground-truth ROI connectivity (bgPDC) in all subjects for $T$ = 60 s. See caption of Supplementary Figure 3 for detail.

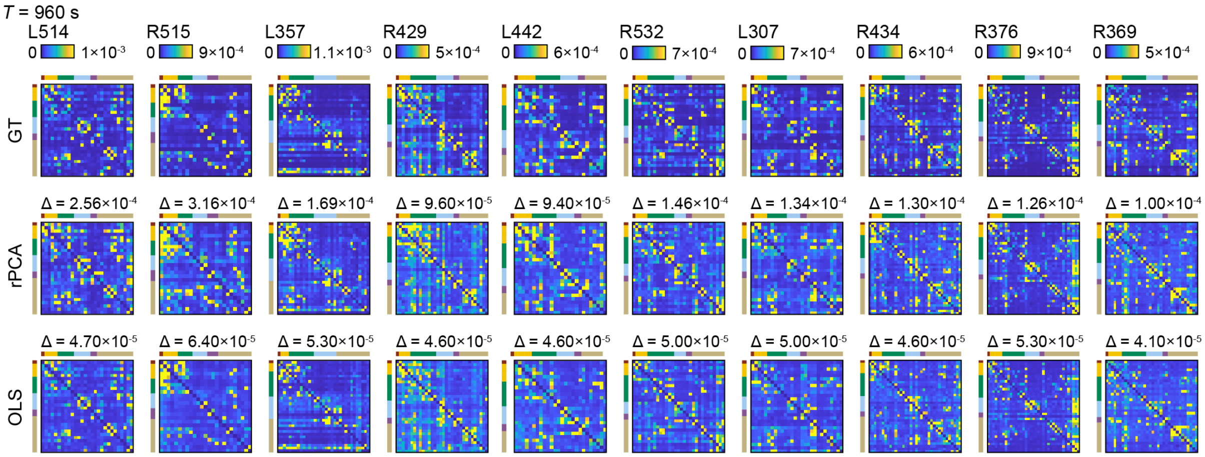


Supplementary Figure 6. Recovery of ground-truth ROI connectivity (bgPDC) in all subjects for $T$ = 960 s. See caption of Supplementary Figure 3 for detail.

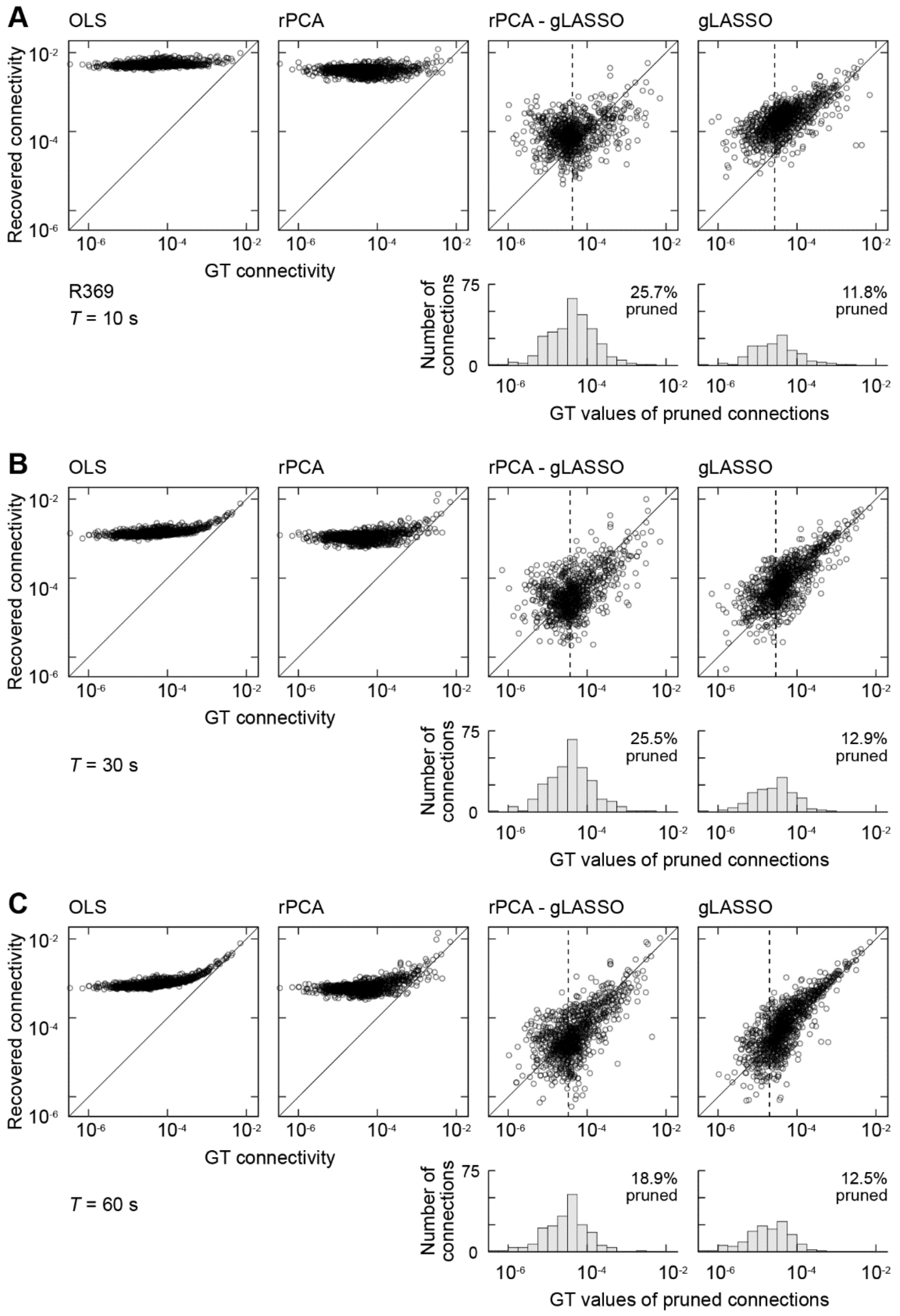


**Supplementary Figure 7.** Comparison of ground-truth vs recovered bgPDC connectivity. Same as Figure 4, but for a different representative subject (R369). See caption of Figure 4 for detail.

### Supplementary Methods

*Invariance of rPCA Method with Invertible Transformations*

The MVAR model of Eq. (1) may be rewritten in terms of the ROI signals $\mathbf{y}_j(n)$, $j=1,2,\ldots,M_R$ as

$$\mathbf{y}_j(n) = \sum_{m=1}^{M_R}\sum_{k=1}^{p} \mathbf{A}_{j,m}(k)\mathbf{y}_m(n-k) + \mathbf{u}_j(n), \quad j = 1,2,\ldots,M_R$$

where $M_R$ is the number of ROIs, $\mathbf{A}_{j,m}(k)$ is the matrix of weights mapping the data in ROI $m$ to ROI $j$ at lag $k$, and $\mathbf{u}_j(n)$ is the vector innovations or model error for ROI $j$. We assume $E\{\mathbf{u}_j(n)\mathbf{u}_j^H(n)\} = \mathbf{\Sigma}_j$. If the PCA transformation $\mathbf{W}_j$ preserves 100% of the variance, then $\mathbf{W}_j$ is a full rank matrix with orthonormal columns, which implies $\mathbf{W}_j^{-1} = \mathbf{W}_j^T$ and the rPCA MVAR model may be expressed as

$$\mathbf{x}_j(n) = \sum_{m=1}^{M_R}\sum_{k=1}^{p} \tilde{\mathbf{A}}_{j,m}(k)\mathbf{x}_m(n-k) + \tilde{\mathbf{u}}_j(n), \quad j = 1,2,\ldots,M_R$$

where $\tilde{\mathbf{A}}_{j,m}(k) = \mathbf{W}_j\mathbf{A}_{j,m}(k)\mathbf{W}_m^T$ and $E\{\tilde{\mathbf{u}}_j(n)\tilde{\mathbf{u}}_j^H(n)\} = \tilde{\mathbf{\Sigma}}_j = \mathbf{W}_j\mathbf{\Sigma}_j\mathbf{W}_j^T$. Note that because of these relationships, we have that

$$\tilde{\bar{\mathbf{A}}}_{j,m}(f) = \mathbf{W}_j\bar{\mathbf{A}}_{j,m}(f)\mathbf{W}_m^T$$

The invertibility of $\mathbf{W}_j$ ensures that connectivity metrics based on $\tilde{\bar{\mathbf{A}}}_{j,m}(f)$ are identical to those based on $\bar{\mathbf{A}}_{j,m}(f)$. For example, consider the term $\bar{\mathbf{A}}_{k,j}^H(f)\mathbf{\Phi}_{kk}\bar{\mathbf{A}}_{k,j}(f)$ that arises in the vector version of the gPDC. We have

$$\begin{aligned}\tilde{\bar{\mathbf{A}}}_{k,j}^H(f)\tilde{\mathbf{\Phi}}_{kk}\tilde{\bar{\mathbf{A}}}_{k,j}(f) &= \left(\mathbf{W}_k\mathbf{A}_{k,j}\mathbf{W}_j^T\right)^H\left(\mathbf{W}_k\mathbf{\Sigma}_k\mathbf{W}_k^T\right)^{-1}\mathbf{W}_k\mathbf{A}_{k,j}\mathbf{W}_j^T \\ &= \mathbf{W}_j\bar{\mathbf{A}}_{k,j}^H(f)\mathbf{\Phi}_{kk}\bar{\mathbf{A}}_{k,j}(f)\mathbf{W}_j^T\end{aligned}$$

Since $\mathbf{W}_j^{-1} = \mathbf{W}_j^T$, the trace and determinant are invariant to the transformation.

If $\mathbf{\Sigma}_j$ is diagonal, then the there is no instantaneous causality within the ROI. Transformation by $\mathbf{W}_j$ will lead to nondiagonal $\tilde{\mathbf{\Sigma}}_j$ in general, indicating that the rPCA approach may introduce instantaneous causality within an ROI. However, the invariance of connectivity to transformations satisfying $\mathbf{W}_j^{-1} = \mathbf{W}_j^T$ indicates that introducing instantaneous causality within an ROI does not compromise connectivity between ROIs.

*ROI-Based Connectivity Metric*

of various covariance matrices associated with the underlying MVAR model. While theoretically appealing, the practical difficulties of reliably computing determinants as the size of the blocks increases in the limited data regime led us to the alternative trace-based measures employed in this paper. Trace based metrics have been considered in lieu of determinant based metrics in other classes of problems, see e.g., (Durieu et al., 1996).

The determinant of a covariance matrix measures volume of the associated error ellipsoid, while the trace measures the sum of the axes (Durieu et al., 1996). This interpretation is consistent with the fact that the determinant of a covariance matrix is the product of the eigenvalues, while the trace is the sum of the eigenvalues. Hence, determinants of even modest-sized covariance matrices are very sensitive to scaling. If the *M*-dimensional matrix ***F*** is replaced by $c\boldsymbol{F}$, the determinant scales by $c^M$, so if $c = 100$ – a factor of ten scaling of the data – and $M=20$, the determinant scales by $10^{40}$. In contrast, the trace scales by *c*, independent of the number of electrodes in the ROI. Geometrically, scaling the axes has a much bigger effect on the volume than on the sum of the axes. Similarly, the determinant is very sensitive to errors in small eigenvalues that arise due to limited data or numerical rounding. Suppose the ordered eigenvalues of a matrix are denoted as $s_1 \geq s_2 \geq \ldots \geq s_M > 0$ and they are estimated with errors $e_1, e_2, \ldots, e_M$. The relative error of the determinant is

$$\frac{\prod_{i=1}^{M} s_i + e_i}{\prod_{i=1}^{M} s_i} = \prod_{i=1}^{M} 1 + \frac{e_i}{s_i}$$

Thus, errors in estimating the smallest eigenvalues have the largest impact, and the relative error can grow in an unbounded manner as $s_M$ shrinks toward zero. As noted in the Discussion, the smallest eigenvalues are the most difficult to estimate reliably from limited data, rendering determinant-based metrics potentially very sensitive to estimation errors associated with limited data. However, the relative sensitivity of the trace is bounded by the size of the error relative to the largest eigenvalue, since

$$\frac{\sum_{i=1}^{M} s_i + e_i}{\sum_{i=1}^{M} s_i} < \sum_{i=1}^{M} 1 + \frac{e_i}{s_1}$$

These observations are consistent with our geometric understanding that a perturbation in one of the smaller dimensions of an ellipsoid can have a very large impact on the volume, whereas the sum of the dimensions is much less affected by a perturbation.

Interpreting the trace of a covariance matrix as the total power associated with the variables, we write the vector version of the squared generalized PDC in (Baccala et al., 2007) capturing the influence of ROI *j* on ROI *i as*

$$\pi_{i,j}^{2^{(Tr)}}(f) = \frac{\mathrm{Trace}\left\{\mathbf{A}_{i,j}^{H}(f)\boldsymbol{\Phi}_{ii}\mathbf{A}_{i,j}(f)\right\}}{M_i \mathrm{Trace}\left\{\sum_{k=1}^{R} \bar{\mathbf{A}}_{k,j}^{H}(f)\boldsymbol{\Phi}_{kk}\bar{\mathbf{A}}_{k,j}(f)\right\}} \tag{S1}$$

We chose to normalize the ratio of the traces by the number of electrodes in region $i$, $M_i$, to facilitate comparison of connectivity to regions with vastly differing numbers of electrodes, but such normalization is not necessary and may not be desirable in other applications. Indeed Eq. (S1) has an upper limit of one without normalization by $M_i$. Clearly Eq. (S1) simplifies to the generalized PDC of (Baccala et al., 2007) in the case of scalar time series as the trace of a scalar is a scalar and $M_i = 1$.

### Data and code availability

Data is available via a request to the Authors pending establishment of a formal data sharing agreement, submission of a formal project outline, and agreement about co-authorship. All software developed for this study will be freely available via Git repository. Please contact Bryan Krause for details.

<u>REFERENCES CITED</u>

Baccala, L.A., Sameshima, K., Takahashi, D.Y., 2007. Generalized partial directed coherence. 15th International Conference on Digital Signal Processing. Ieee, Cardiff Univ, Cardiff, WALES, pp. 163-+.

Durieu, C., Polyak, B.T., Walter, E., 1996. Trace Versus Determinant in Ellipsoidal Outer-Bounding, with Application to State Estimation. IFAC Proceedings Volumes 29, 3975-3980.

Faes, L., Nollo, G., 2013. Measuring frequency domain granger causality for multiple blocks of interacting time series. Biol Cybern 107, 217-232.

Geweke, J., 1982. Measurement of Linear-Dependence and Feedback between Multiple Time-Series. Journal of the American Statistical Association 77, 304-313.

Pordes, R., Petravick, D., Kramer, B., Olson, D., Livny, M., Roy, A., Avery, P., Blackburn, K., Wenaus, T., Wuerthwein, F., Foster, I., Gardner, R., Wilde, M., Blatecky, A., Mcgee, J., Quick, R., 2007. The open science grid. Scidac 2007: Scientific Discovery through Advanced Computing 78.
